# Differential Roles of Kinetic On- and Off-Rates in T-Cell Receptor Signal Integration Revealed with a Modified Fab’-DNA Ligand

**DOI:** 10.1101/2024.04.01.587588

**Authors:** Kiera B. Wilhelm, Anand Vissa, Jay T. Groves

## Abstract

Antibody-derived T-cell receptor (TCR) agonists are commonly used to activate T cells. While antibodies can trigger TCRs regardless of clonotype, they bypass native T cell signal integration mechanisms that rely on monovalent, membrane-associated, and relatively weakly-binding ligand in the context of cellular adhesion. Commonly used antibodies and their derivatives bind much more strongly than native peptide-MHC (pMHC) ligands bind their cognate TCRs. Because ligand dwell time is a critical parameter that tightly correlates with physiological function of the TCR signaling system, there is a general need, both in research and therapeutics, for universal TCR ligands with controlled kinetic binding parameters. To this end, we have introduced point mutations into recombinantly expressed α-TCRβ H57 Fab to modulate the dwell time of monovalent Fab binding to TCR. When tethered to a supported lipid bilayer via DNA complementation, these monovalent Fab’-DNA ligands activate T cells with potencies well-correlated with their TCR binding dwell time. Single-molecule tracking studies in live T cells reveal that individual binding events between Fab’-DNA ligands and TCRs elicit local signaling responses closely resembling native pMHC. The unique combination of high on- and off-rate of the H57 R97L mutant enables direct observations of cooperative interplay between ligand binding and TCR-proximal condensation of the linker for activation of T cells (LAT), which is not readily visualized with pMHC. This work provides insights into how T cells integrate kinetic information from synthetic ligands and introduces a method to develop affinity panels for polyclonal T cells, such as cells from a human patient.

**STATEMENT OF SIGNIFICANCE:** T cells read kinetic information from ligands binding to T-cell receptors (TCRs) to make cell fate decisions. Unique kinetic features of a modified Fab’-DNA ligand enable direct visualization multiple TCR signal coordination through a nascent LAT condensation event. We further observed positive feedback through a kinetic on-rate enhancement in the growing LAT condensate. These observations help unify several seemingly disparate aspects of TCR signaling that have been debated in the literature. Furthermore, calibration of the Fab’-DNA ligand against native agonist pMHC establishes a basis for quantitative analysis of TCR signaling in polyclonal T cell populations.

## INTRODUCTION

T cells integrate kinetic information from the binding interaction between peptide major histocompatibility complex (pMHC) ligands and T-cell receptors (TCRs) to make cell fate decisions. Developing thymocytes undergo differential positive or negative selection across a narrow affinity range (1, 2), while T cells in the periphery receive survival signals from low-affinity self-pMHC and activate and differentiate in response to higher-affinity pathogenic pMHC (3). Ligand potency is well-known to correlate with binding kinetics, with high affinity and long-dwelling ligands stimulating greater T cell activation in numerous studies (4–13). However, while very strong TCR binders are the most potent in *in vitro* assays, they do not necessarily elicit the greatest immune response *in vivo* (14). Affinities for native pMHC binding TCR range from 1-10 µM for agonist pMHC (2-10 s dwell time) and > 10 µM for self or null pMHC (< 0.5 s dwell time), which is quite weak for ligand–receptor interactions. The T cell signaling network is naturally tuned to respond to TCR ligands in the low-to mid-micromolar range (15–17).

Mechanistic understanding of how T cells integrate information from binding kinetics, including both native and non-physiological TCR binding parameters, is of critical importance given the rapid development of synthetic T cell engagers for cancer immunotherapy. Current efforts to therapeutically direct cytotoxic T cell activity to cancerous cells typically employ moieties that bind receptors on T cells very strongly. Bispecific T cell engagers use antibody-derived fragments with low-nanomolar affinity to bridge tumor markers to TCRs (18, 19). While a few T cell engagers have been approved for clinical use, on-target, off-tumor effects from the directed cytotoxic T cell activity often cause serious clinical complications such as cytokine storm (20–22). Alternatively, chimeric antigen receptors (CARs) with antibody-derived tumor-binding moieties have been introduced into patients’ T cells to redirect cellular activity to the target of interest. Some CAR therapies have shown impressive clinical success; however CARs fail to trigger classic signatures of T cell activation (23), have blunted antigen sensitivity (24), and are similarly accompanied by toxic on-target, off-tumor effects (25). There is growing acknowledgement that ligand–receptor binding affinity of therapeutics must be carefully considered in order to maximize efficacy while minimizing adverse side effects (26–29). The specific details of how to optimally tune binding kinetics remains under intensive investigation, in part because the mechanism by which T cells read and integrate TCR binding kinetics remains substantially unresolved.

We have previously introduced a class of antibody-derived T cell engagers to universally activate T cells in quantitative supported lipid bilayer (SLB) assays (30). These Fab’-DNA ligands — built from anti-TCR/CD3 Fab’ fragment covalently linked to a short DNA oligonucleotide — mimic many properties of physiological TCR triggering and cellular activation by pMHC ligands. They are strictly monovalent, and like native agonist pMHC they are incapable of activating T cells from solution at any concentration (31), but become potent activators when bound to a membrane surface (30, 32–34). Single ligation events between Fab’-DNA and TCR are sufficient to produce significant TCR-proximal phosphorylation activity (35), and cells can activate in response to just a handful widely-spaced ligation events (10, 30). The anti-TCR/CD3 Fab’ fragments bind constant regions of the TCR complex rather than the hypervariable pMHC recognition site, and thus they are able to activate T cells regardless of TCR clonotype. Like other antibody-based TCR binders, the Fab’-DNA ligands developed thus far have very high affinities and long dwell times, binding the TCR for a mean dwell time of well over 2 min compared to a range of subsecond to about 10 seconds for self- and agonist-pMHC.

Here, we express the H57-597 α-TCRβ Fab fragment (henceforth H57 Fab) recombinantly in *E. coli* and modulate its affinity by introducing point mutations at the Fab–TCR binding interface (Fig. 1). Single molecule fluorescence imaging provides direct measurements of Fab’-DNA– TCR binding interactions at the junction between an SLB and a primary murine CD4^+^ effector T cell. The R97L mutant disrupts two hydrogen bonds at the binding interface, resulting in a 4 s mean dwell time — similar to native pMHC agonists (34) — while retaining a high on-rate. We investigate cellular activation in response to H57 R97L Fab’-DNA by imaging translocation of the nuclear factor of activated T cells (NFAT) from the cytosol to the nucleus, which provides a robust readout of signaling through the calcium pathway (36). The dose-response curve of cells clonally expressing the AND TCR in response to H57 R97L Fab’-DNA falls between curves for the long-dwelling MHC class II I-E^k^ loaded with moth cytochrome c (MCC) peptide (I-E^k^/MCC) and shorter-dwelling I-E^k^/T102S. Moreover, parental H57 Fab’-DNA and the R97L mutant are able to activate polyclonal murine effector T cells, with potency correlating with dwell time.

**Figure 1.**
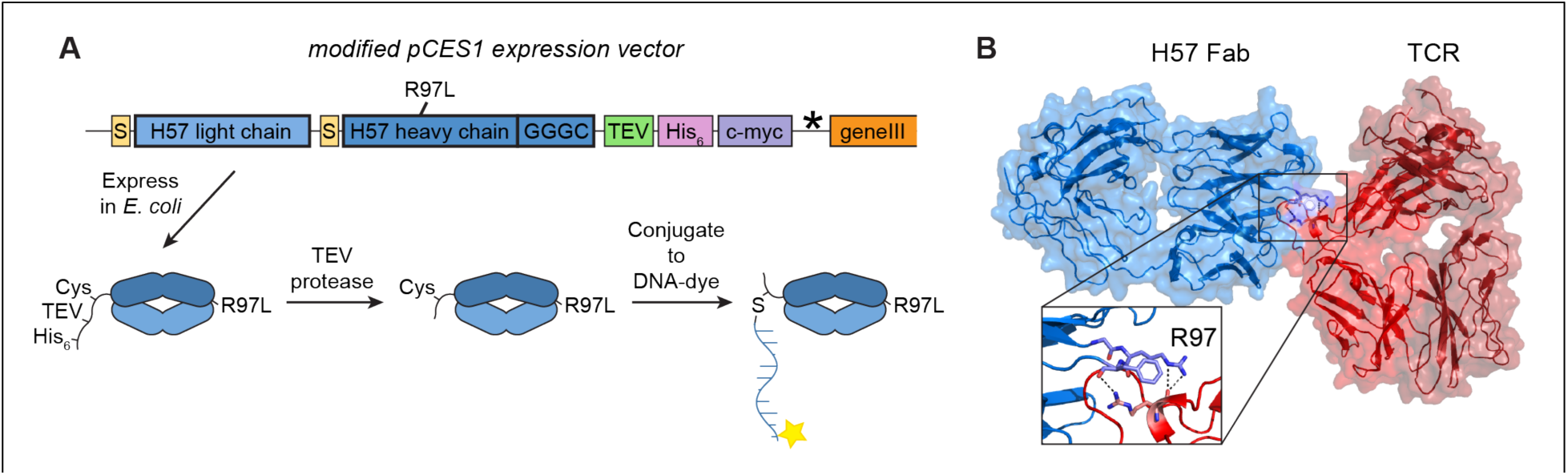
H57 Fab’ expression enables the introduction of affinity-modulating mutations. (A) The pCES1 expression vector was used to express H57 Fab in *E. coli*. An R97L mutation was introduced in the heavy chain to decrease the affinity between the H57 Fab and TCR. A GGGC linker allowed for conjugation to a dye-labeled DNA oligonucleotide. (B) The crystal structure of H57 Fab bound to TCR shows hydrogen bonding contacts between R97 and F98 on H57 Fab heavy chain and R120 on TCRβ. PDB: 1NFD.

We further examine TCR-proximal signaling by imaging the linker for activation of T cells (LAT), which undergoes a phosphorylation-driven condensation (37, 38) in response to a single ligated TCR (13). Using these single-molecule imaging methods, we find that single Fab’-DNA–TCR binding events have the same propensity to form LAT condensates as pMHC (13), suggesting that signal from TCR ligation is initially integrated by the same cellular machinery in these CD4^+^ effector T cells despite Fab’-DNA not engaging co-receptor. We speculate that co-receptor effects would be more pronounced in thymocytes, in CD8+ cells, and/or for shorter-dwelling ligands ( < 2 s) (39–41). Though ligand clustering is not a requirement for cellular activation (32, 33, 42), the unique binding kinetics of H57 R97L Fab’-DNA allow us to directly image how cooperativity enhances activity for short-dwelling ligands by allowing multiple triggered TCRs to contribute to the same local pool of phosphorylated LAT. Such cooperativity mechanisms involving multiple TCR and weak pMHC ligands have long been speculated to exist (9, 15, 43–46), but their direct observation has proven elusive. This imaging study provides mechanistic insight into how ligand binding kinetics are read and integrated by the T cell signaling network, particularly at early stages, and builds a framework for developing synthetic anti-TCR ligands with binding kinetics more similar to pMHC to quantitatively study signal integration in polyclonal T cell populations.

## RESULTS

### H57 Fab’ expression enables the introduction of affinity-modulating mutations

Commercial α-TCR antibodies have been developed to bind TCR with high affinities. We adopted the pCES1 Fab expression vector (47, 48), to express the H57 Fab in *E. coli* and introduced point mutations that decrease binding affinity to more closely mimic pMHC (Fig. 1A). A GGGC linker was appended directly to the heavy chain for conjugation to maleimide-functionalized DNA and a TEV recognition site was introduced to cleave off the His- and myc-tags after purification. An amber stop codon separated the tags and geneIII, terminating expression after the heavy chain in standard *E. coli* strains and enabling phage display in amber suppressor strains. While phage display was not used in this study, it could readily be used to generate a library of Fab ligands with diverse binding kinetics.

Informed by its crystal structure (49), targeted point mutations were introduced into H57 Fab to weaken interfacial contacts. The complementarity-determining region 3 of the H57 Fab heavy chain forms numerous atomic contacts with TCRβ, including hydrogen bonding between R120 on TCRβ and R97 and F98 on the Fab (Fig. 1B). An Arg to Leu mutation that decreases affinity in the anti-human CD3ε OKT3 Fab fragment by breaking a hydrogen bond (50) suggested that a similar mutation in H57 could be effective. Purifying the H57 R97L Fab’ in *E. coli* and conjugating to a dye-labeled DNA oligonucleotide through maleimide-cysteine chemistry (30) yielded a clean sample of H57 R97L Fab’-DNA for analysis in live-cell imaging assays (Fig. S1).

### R97L Fab’ mutant has a pMHC-like off-rate but with a high on-rate that enhances rebinding

Binding kinetics between H57 R97L Fab’-DNA and TCR were characterized at the interface between SLBs and primary AND CD4^+^ effector T cells in the context of integrin adhesion at 37 °C (Fig. 2A). Primary murine lymphocytes and spleenocytes were expanded with MCC peptide and imaged 5-8 days after harvesting (details in Materials and Methods). Intercellular adhesion molecule 1 (ICAM-1) binding to the integrin receptor lymphocyte function-associated antigen 1 (LFA-1) stably adhered T cells to the SLB with large cell footprints and enabled native crosstalk between triggered TCR and LFA-1 (51). Fluid, low-defect SLBs were formed from 95% DOPC, 3% MCC-PE, and 2% Ni-NTA-DOGS lipids. A thiol-functionalized DNA oligonucleotide, complementary to the Fab’-DNA, was covalently linked to MCC-PE lipids at densities of 100’s µm^-2^, and Fab’-DNA density on the bilayer was precisely controlled to range from 0.05 – 10 µm^-2^ by varying its incubation concentration and duration. ICAM-1 coupled to SLBs at 100’s µm^-2^ through multivalent interactions between the His-tag on ICAM-1 and Ni^2+^ chelated by NTA-DOGS lipids.

**Figure 2.**
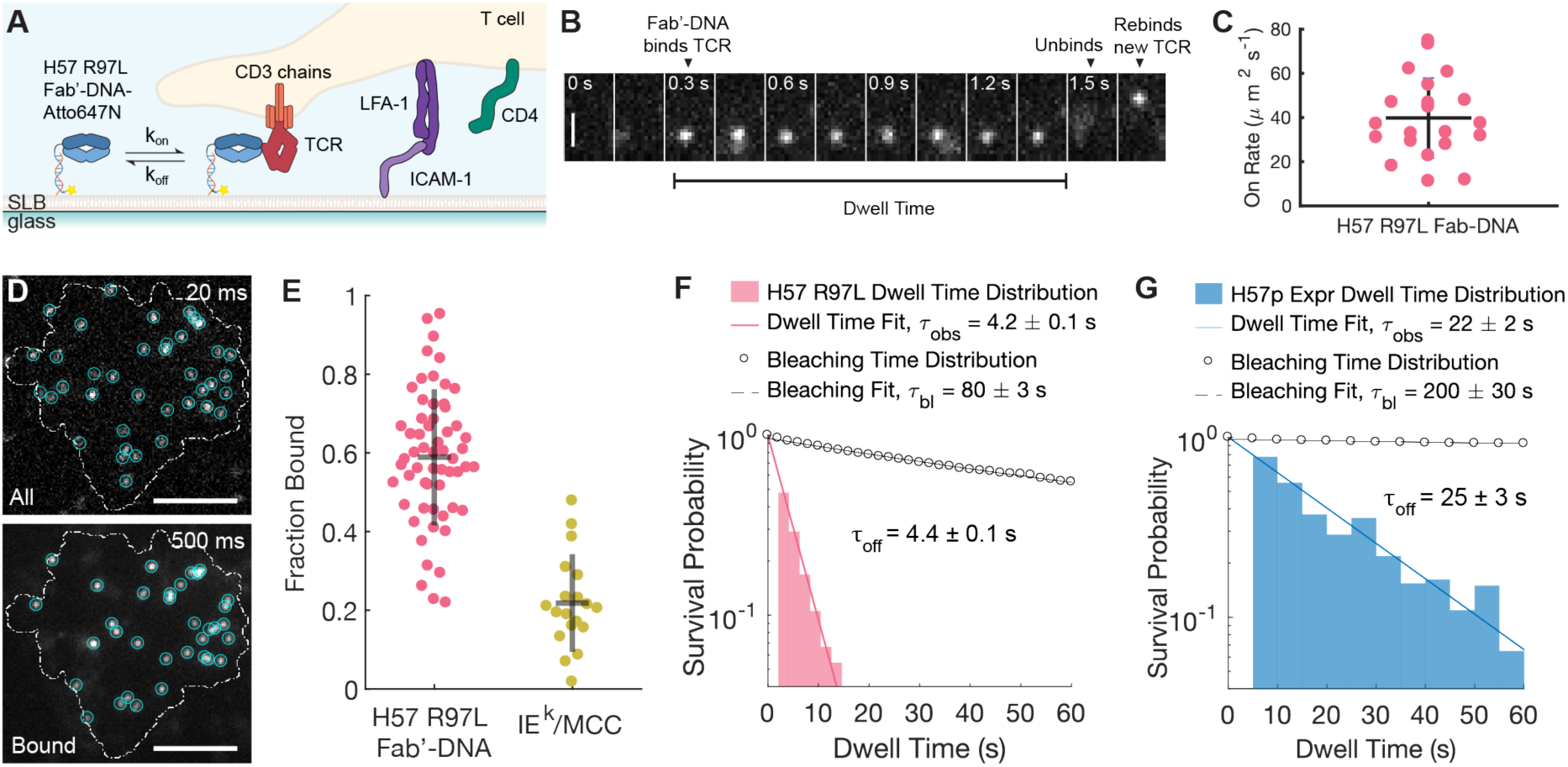
R97L Fab’ mutant has a pMHC-like off-rate but with a high on-rate that enhances rebinding. (A) H57 R97L Fab’-DNA-Atto647N presented on a glass-supported lipid bilayer binds to TCR in the context of LFA-1:ICAM-1 adhesion in live cell assays. (B) A single molecule of Fab’-DNA binds TCR (0.3 s) forming a well-resolved image with TIRF microscopy. The molecule unbinds at 1.5 s, forming a blurred image, and rebinds about a micron away shortly thereafter. The dwell time of a single binding event is directly visualized. Scale bar 1 µm. (C) The measured cellular on-rate of H57 R97L Fab’-DNA is 40 ± 20 µm^2^s^-1^ (SD). n = 21 cells, 2 mice. (D) All H57 R97L Fab’-DNA under a cell footprint are imaged in TIRF using a short exposure time, and bound ligand are resolved using a long exposure time. (E) The fraction of ligand bound is 0.6 ± 0.2 (SD) on bilayers presenting 0.1 µm^-2^ H57 R97L Fab’-DNA. n = 59 cells, 1 mouse. On bilayers presenting 0.48 µm^-2^ IEk/MCC, the fraction bound is 0.2 ± 0.1 (SD). n = 19 cells, 1 mouse. (F) The observed dwell time distribution of H57 R97L Fab’-DNA fits a single exponential decay with an average dwell time of 4.2 ± 0.1 s (95% CI). The dwell time is 4.4 ± 0.1 s (95% CI) after correction for photobleaching n = 6 cells, 2 mice, 759 dwell events. (G) The observed dwell time distribution of expressed, parental H57 Fab’-DNA fits a single exponential decay with an average dwell time of 22 ± 2 s (95% CI). The dwell time is 25 ± 3 s (95% CI) after correction for photobleaching n = 7 cells, 2 mice, 994 dwell events.

Total internal reflection fluorescence (TIRF) microscopy of the live cell-SLB interface provides robust imaging capabilities, down to the single molecule level, of receptor binding and membrane-proximal signaling reactions. Single binding interactions between Fab’-DNA and TCR were directly observed by continuously imaging Fab’-DNA-Atto647N with a 50 ms exposure time (Movie S1). While unbound Fab’-DNA-Atto647N underwent Brownian motion with a diffusion coefficient of 2-3 µm^-2^ (30), Fab’-DNA molecules bound to TCR had dramatically reduced mobility. A montage of a selected region under a T cell shows the image of a Fab’-DNA molecule becoming clearly resolved upon TCR binding (0.3 s), blurring upon unbinding as the ligand quickly diffuses (1.5 s) and becoming clear again upon rebinding a new TCR (1.65 s) (Fig. 2B).

Rebinding has been much discussed in the literature as a mechanism for ligand discrimination and signal enhancement (7–9, 44), but to our knowledge, it has not previously been directly observed by single molecule imaging. The unique combination of high on-rate and short dwell time enabled direct visualization of single H57 R97L Fab’-DNA molecules unbinding from one TCR and rebinding to a different TCR, which occurred quite commonly. Two Fab’-DNA molecules show this rebinding behavior in the same cell within a couple seconds (Box 1 and 2 of Movie S1). Rebinding is not seen with IE^k^/MCC pMHC in analogous measurements on supported membranes (10) and was determined to be exceedingly unlikely for typical pMHC molecules due to their intrinsically low kinetic on-rates for TCR binding (34). *In vivo,* where other features of the cell-cell interface may effectively corral pMHC and/or TCR, it is plausible that rebinding of pMHCs to multiple TCRs occurs more readily despite intrinsically slow on-rates (7).

Taking advantage of the dramatic change in mobility upon ligand binding, we used a long exposure times (500 ms) at low excitation powers (0.3 mW) to clearly resolve and track individual Fab’-DNA–TCR complexes while free Fab’-DNA appeared as a diffuse background (Fig. S2A) (10, 34). We directly quantitated *in situ* kinetic on- and off-rates by tracking the rate of new binding interactions and duration of binding events, respectively. The H57 R97L Fab’-DNA–TCR on-rate was measured to be 40 ± 20 µm^2^s^-1^ (Fig. 2C), ten-fold greater than the on-rate measured between IE^k^/MCC and the AND TCR (13). The fraction of ligand bound to TCR under a cell footprint was used to measure the efficiency of ligand-receptor binding. All (Fig. 2D, top, 20 ms exposure) and bound (Fig. 2D, bottom, 500 ms exposure) Fab’-DNA were imaged sequentially in TIRF and the T cell footprint was visualized using reflection interference contrast microscopy (RICM). On bilayers presenting 0.1 µm^-2^ H57 R97L Fab’-DNA, a high fraction, 0.6 ± 0.2, of Fab’-DNA ligands under cell footprints were bound to TCR (Fig. 2E). This is not as high as Fab’-DNA constructs derived from commercially available ligands, for which nearly all ligands under a cell are bound (30, 35), but is significantly higher than fraction bound measured for IE^k^/MCC pMHC (Fig. 2E), in agreement with the measured on-rates.

With Fab’-DNA presented at low density on the SLB (0.08 µm^-2^), single ligation events between Fab’-DNA and TCR are spaced microns apart and often remain distinct for their entire lifetime, facilitating tracking and analysis of single events (Fig. S2A). Binding events were identified by a single step appearance in fluorescence intensity compared to background and tracked until bleaching or unbinding in a single step (Fig. S2B). The duration of every binding event was compiled into a dwell time distribution for H57 R97L Fab’-DNA, and the distribution was well-fit by a single exponential decay, indicating first-order unbinding kinetics (Fig. 2F). After correcting for photobleaching, the mean dwell time for H57 R97L Fab’-DNA was determined to be 4.4 ± 0.1 s. This is much shorter than the >2 min dwell time of H57 Fab’-DNA synthesized from commercially available antibody (30) and similar to the 5.2 s mean dwell time of native 5c.c7 TCR binding to its native agonist IE^k^/MCC pMHC (34).

Fab’-DNA synthesized with parental H57 Fab’ expressed in *E. coli* exhibited a mean dwell time of 25 ± 3 s (Fig. 2G). The reduced dwell time compared to commercially available antibody may be attributed to the expressed Fabs lacking glycosylation. As the R97L mutant well-matched the dwell time of native pMHC–TCR interactions, this construct was the focus of further experiments.

### Single binding events between H57 R97L Fab’-DNA and TCR lead to LAT condensation

We next investigated how local signaling from isolated H57 R97L Fab’-DNA–TCR complexes compared to that from pMHC ligands through detailed analyses of multicolor TIRF imaging of bound Fab’-DNA-Atto647N at the SLB-T cell interface and LAT-eGFP in the T cell plasma membrane (Movie S2). Phosphorylated LAT condenses in response to individual TCR triggering events and serves as a critical gating mechanism for signal propagation (13). A bicistronic plasmid containing LAT-eGFP and NFAT-mCherry, separated by a self-cleaving P2A peptide (52), was transduced into T cells two days after organ harvest using murine stem cell virus retroviral transduction. Overexpression of fluorescently-tagged LAT does not change signaling outcomes in T cells (13), and the NFAT reporter lacks a DNA-binding domain, so it also does not affect downstream signaling (36).

A representative time-sequence of images shows a single Fab’-DNA that bound TCR (0:02 time point) and subsequently triggered the formation of a local LAT condensate (0:12) (Fig. 3A). The bound Fab and LAT condensate moved together for their lifetimes, though the center of the LAT condensate drifted somewhat from the location of the bound TCR, indicating weak physical coupling (Fig. 3B). Significant signal amplification occurs upon LAT condensation, with one binding event leading to hundreds of phosphorylated LAT molecules in the condensate (Fig. 3C) (13). LAT condenses together with, and is cross-linked by, adapter proteins like growth factor receptor-bound 2 proteins (Grb2) and critical enzymes such as Son of Sevenless (SOS) and phospholipase C γ1 (PLCγ1) (53).

**Figure 3.**
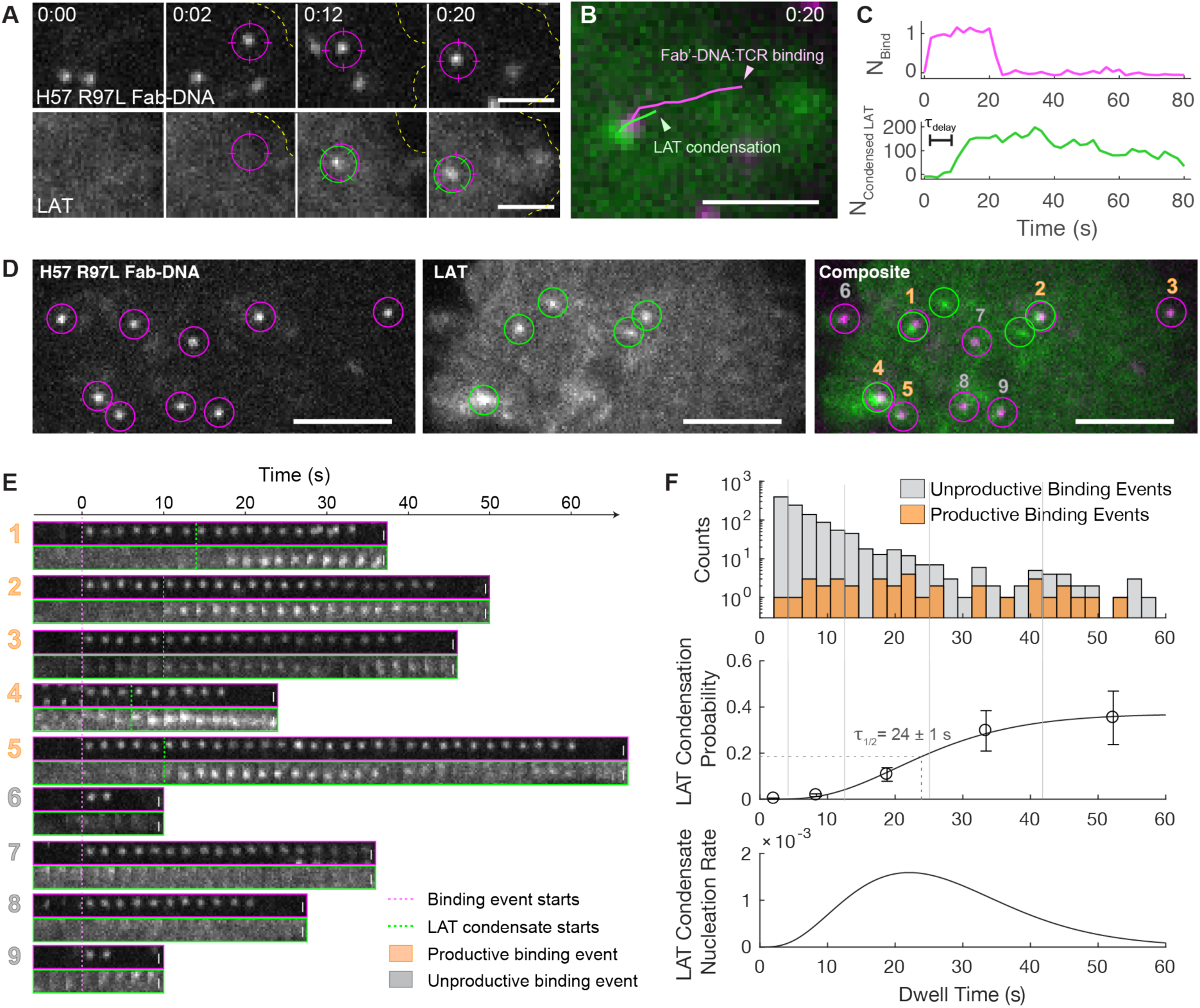
Long, single binding events between H57 R97L Fab’-DNA and TCR lead to LAT condensation. (A) A representative montage shows a single binding event (top, magenta) followed by formation of a proximal LAT condensate (bottom, green). Scale bar 2 µm. (B) The binding event and LAT condensate stay colocalized through time. Scale bar 2 µm. (C) This single binding event causes condensation of 100’s of LAT molecules. (D) In a stillframe, several binding events colocalize with LAT condensates. Scale bar 5 µm. (E) All binding events that appear in (D) (top) and the local LAT signal (bottom) are tracked through time. Pink and green dashed lines note initial Fab’-DNA binding and LAT condensation, respectively. Scale bars 0.5 µm. Imaging data are representative of 15 cells from 5 mice. (F) (Top) The dwell time distributions of productive binding events is divided by that of all binding events to (Middle) calculate the LAT condensation probability given a binding event dwell time. n = 6 cells, 2 mice. Error bars note SEM. Solid grey vertical lines mark bins used to calculate LAT condensate probability. Data are fit with a regularized gamma function (black line). The inflection point in the fit is τ1/2 = 24 ± 1 s (95% CI) (dotted grey line). (Bottom) The fit is used to calculate the rate of LAT condensate nucleation, also known as the LAT propensity function.

Several single Fab’-DNA–TCR complexes and associated LAT condensates were typically observed over the T cell–SLB interface (Fig. 3D). At the low H57 R97L Fab’-DNA density of 0.08 µm^-2^, most LAT condensates colocalize with a single binding event. Montages of these binding events through time illustrate the relative timing of binding and LAT condensation (Fig. 3E), and exhibit striking similarity to analogous data collected with pMHC ligands (13). The first five montages show H57 R97L Fab’-DNA binding TCR (top) followed by proximal LAT condensation after a significant delay of several seconds (bottom). Montages six through nine show binding events for which LAT never condenses locally. Notably, these unproductive binding events are among the shortest of those that appear in the snapshot in Fig. 3D.

Single binding events that remained isolated for the duration of their visible lifetime were categorized as either productive (produced local LAT condensate) or unproductive (did not) and histogrammed by their dwell time (Fig. 3F, top). Binning the data by Fab’-DNA dwell time (vertical gray lines) and dividing the number of productive binding events by total number of events yielded the probability of LAT condensation given a dwell time (Fig. 3F, middle). These data represent the measurement of a single TCR antigen discrimination function, i.e. the dwell-time-dependent probability that a single ligated TCR contributes to the cell-wide activation response. The dwell time at which binding events have half-maximal probability of generating a local LAT condensate is 24 ± 1 s. This time is about 5 times longer than the mean H57 R97L Fab’-DNA:TCR dwell time of 4.4 ± 0.1 s. Moreover, this LAT condensation probability function is nearly identical to that of IE^k^/MCC and IE^k^/T102S pMHCs (13), providing strong evidence that cells integrate signal from Fab’-DNA and pMHC using similar cellular machinery and employ a similar kinetic proofreading mechanism at the level of LAT condensation. Differential effects of co-receptor scanning (41) appear inconsequential at these time scales (pMHC–TCR dwell time greater than a couple of seconds) with effector T cells in supported bilayer assays.

The probability of a local LAT condensate nucleating at a given time interval after ligand binding, known in stochastic kinetics as the propensity function (54), is shown in Fig. 3F, bottom. This function was obtained from the regularized gamma function fit to the probability of LAT condensation as described in McAffee et al. (13). Like with pMHC ligands, the curved rise of the LAT propensity function suggests ≥ 2 kinetic proofreading steps occur between ligand binding and LAT condensation. The fall of the curve back to zero at later time delays indicates that a bound TCR can only produce a LAT condensate within a first minute of its lifetime, and additional dwelling beyond that time has a near-zero probability of being productive.

The delay time between initial H57 R97L Fab’-DNA binding and LAT condensation (τ_delay_) is a direct experimental measure of the LAT propensity function convoluted with ligand photobleaching (Fig. S3A). The histogram of delay times has a mean delay of 15.6 s, nearly four times as long as the mean ligand dwell time before correcting for photobleaching (Fig. 2F). As it is only possible to extract a delay time from binding events that haven’t bleached before LAT condensation commences, the measured delay time is an underestimate. These data emphasize that, when isolated, rare, long-dwelling binding events at the tail end of the dwell time distribution are primarily responsible for generating significant downstream signaling activity. The kinetic off-rate sets the likelihood that a given single ligand dwells long enough to produce a LAT condensate.

### Ligand binding and LAT condensation cooperatively enhance each other

The high on- and off-rates of H57 R97L Fab’-DNA increase the likelihood (compared to low-on-rate pMHCs or long-dwelling Fabs) that short ligation events occur nearby one another in space and time when presented on SLBs at densities where individual binding interactions are countable in TIRF microscopy. Representative time series from cells on bilayers with moderate H57 R97L Fab’-DNA density (0.1–0.2 µm^-2^) illustrate how T cells integrate information across multiple spatiotemporally correlated bound TCR.

First, existing LAT condensates grow and are more persistent when additional Fab’-DNA ligands bind TCR within or near the condensate (Fig. 4A-C; Movie S3). In a representative time series, a single H57 R97L Fab’-DNA–TCR binding event (0:06) initiates a LAT condensate (0:16). Additional ligands bind TCR in the diffraction-limited area (0:28) and the LAT condensate subsequently grows (0:40). The intensity trace of the Fab’-DNA channel shows stepwise, integer changes in fluorescence intensity, as new ligands bind into the area and unbind or photobleach (Fig. 4B). The intensity of the LAT condensate experiences bursts of growth at time delays after new ligand binding (Fig. 4C). The time series shown in Fig. S4A-B (Movie S4) illustrates the differing fates of LAT condensates that experience multiple (Event I) versus no (Event II) additional Fab’-DNA–TCR binding events after LAT condenses. In this montage, both the top (Event I) and the bottom (Event II) Fab’-DNA–TCR complexes initiate a LAT condensate. In Event I, another Fab’-DNA binds into the same diffraction-limited spot, and the condensate substantially grows and persists for about 40 s. In contrast, no further Fab’-DNA ligands bind into the condensate from Event II, the LAT condensate remains small, and it only persists for about 15 s. These examples demonstrate that spatiotemporally-correlated binding enhances local LAT condensation, and suggest a mechanism by which ligands with high on-rates may be more potent than would be assumed by their dwell time alone.

**Figure 4.**
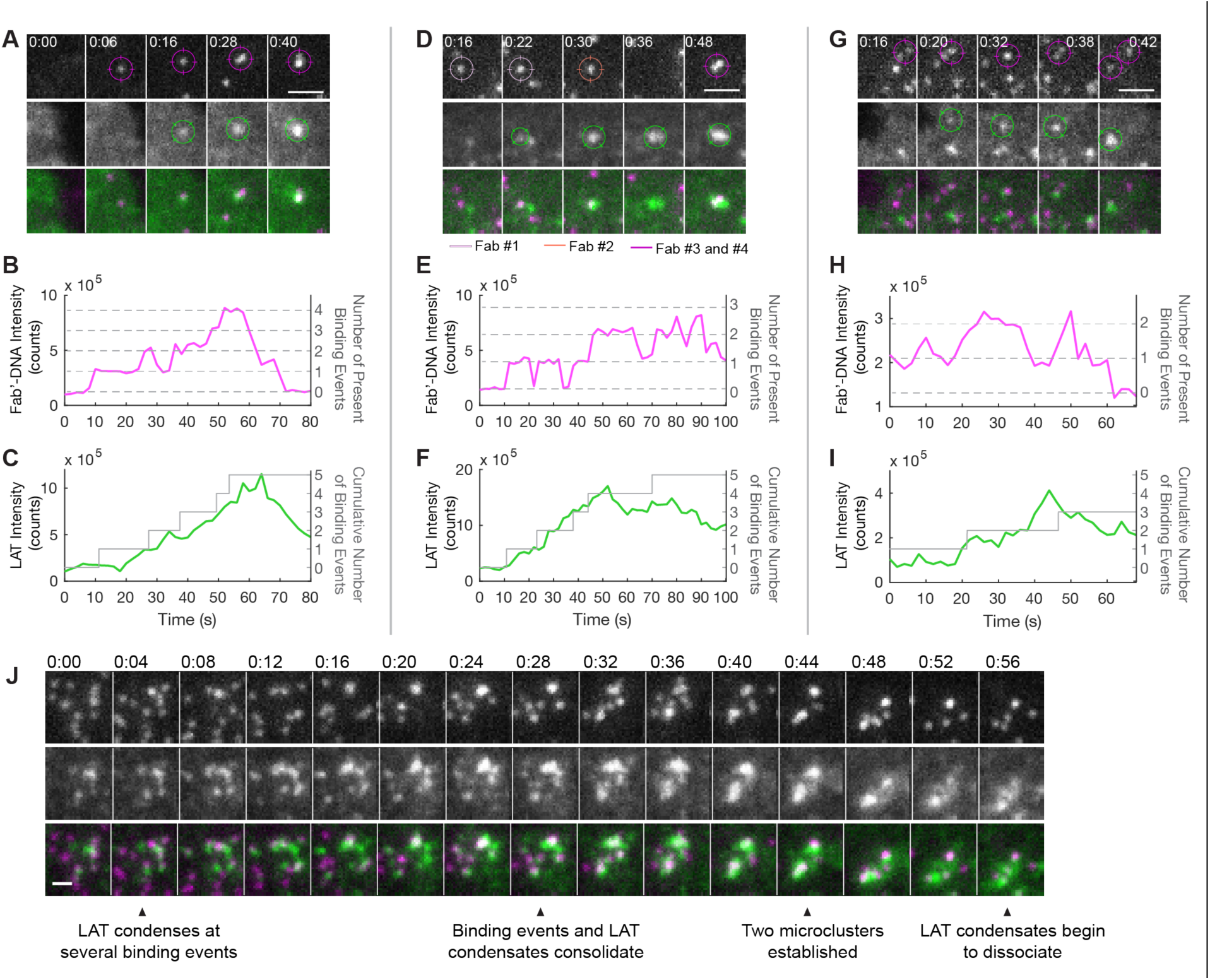
Ligand binding and LAT condensation cooperatively enhance each other. (A) A single H57 R97L Fab’-DNA–TCR binding event (0:06) initiates a LAT condensate (0:16). Additional binding events join in a diffraction limited area (0:28) and the LAT condensate grows (0:40). (B) The intensity trace of the Fab’-DNA-Atto647N signal in (A) shows stepwise, integer increases and decreases in fluorescence intensity, indicating binding directly into the local area and photobleaching or unbinding, respectively. (C) The intensity of the LAT condensate in (A) increases, with a delay, in response to new binding events. The cumulative number of Fab’-DNA that have bound into the tracked area is indicated in gray. (D) A binding event (0:16) initiates a LAT condensate (0:22), which merges with an existing neighboring condensate (0:30). Fab’-DNA molecules unbind from the area or bleach, while the LAT condensate persists (0:36). More Fab’-DNA bind into the area of the condensate, and the condensate grows (0:48). (E) The Fab’-DNA intensity trace shows discrete binding and unbinding/photobleaching events. Varied Fab’-DNA intensity at the end of the trace is due to some persistent binding events drifting into and out of the area of the LAT condensate. (F) The intensity of the LAT condensate increases, with a delay, in response to the first two binding events, in particular. The condensate appears to have a more muted response to later binding events. (G) A LAT condensate begins localized at a single binding event (0:20) and a nearby binding event merges into the same diffraction-limited area shortly thereafter (0:32). One binding event drifts away (0:38) and the other tracks with the growing condensate (0:42). (H) The Fab’-DNA fluorescence trace is more continuous rather than stepwise, indicating fluorophores are moving into and out of the area of the condensate. (I) The LAT intensity trace shows the condensate initiates with a single binding event but grows after the second joins. Scale bars 2 µm. (J) An exemplar montage the interplay between binding events and LAT condensation upon cell landing. Initially, binding events are independent and well-resolved. Some initiate LAT condensation. Binding events merge and LAT condensates coalesce through time, leading to the formation of colocalized Fab’-DNA and LAT microclusters. Scale bar 2 µm.

Second, LAT condensation, in turn, promotes local ligand binding. The representative time series presented and analyzed in Fig. 4 D-F shows numerous independent Fab’-DNA ligands binding to and unbinding from TCR in a condensate (Movie S5). Condensate size increases with the cumulative number of new local binding events rather than sensing the instantaneous number of locally bound Fab’-DNA or the individual persistence of those binding events (Fig. 4F). Like in the Fig. 4A, multiple Fab’-DNA ligands bind directly into the area of the condensate, suggesting that LAT condensates contain a high density of ligand-accessible TCR. In each of these two examples, five new binding events occur within one minute and 0.5 µm^-2^. With the measured cell-wide H57 R97L Fab’-DNA on-rate of 40 ± 20 (SD) µm^2^s^-1^, ligand density of 0.1 µm^-2^ and a cell footprint of 100 ± 25 (SD) µm^2^, 1 ± 0.5 (SD) binding events are expected to occur on average per 0.5 µm^-2^ of contact area within 60 s (see Materials and Methods for calculation details). Within these two LAT condensates, the rate of 5 new visible binding events in this spatiotemporal window is more than 8 standard deviations above the mean.

Observations of a single Fab’-DNA hopping between TCR in a condensate provides further evidence that LAT condensates contain a high density of accessible TCR. An exemplar time series shows a single Fab’-DNA taking multiple unusually large steps between adjacent frames while remaining within a LAT condensate (Fig. S4C) (Movie S6). The frequency of these large steps is about three times greater than the frequency predicted by the step size distribution of bound Fab’-DNA with the same image acquisition parameters (Fig. S4D), suggesting that at least some of these large steps can be attributed to the Fab’-DNA unbinding one TCR and quickly rebinding a nearby TCR, similar to the behavior seen in Movie S1. The apparent increased on-rate within LAT condensates suggests a positive feedback mechanism in which more binding events strengthen a local LAT condensate, which in turn increases the local density of accessible TCR to promote further binding.

Third, well-spaced, bound TCRs consolidate to a diffraction-limited spot within LAT condensates, and this consolidation increases LAT condensate size and lifetime. In the representative time series presented and analyzed in Fig. 4 G-I (Movie S7), LAT begins to condense proximal to a single bound Fab’-DNA (0:20, top right Fab’-DNA). Seconds later, a nearby Fab’-DNA–TCR complex merges into the same diffraction-limited area (0:32). After merging, the LAT condensates strengthen (0:38), and the small cluster of bound Fab’-DNA and LAT condensates remain briefly colocalized. After some time, one of the bound TCR decouples from the condensate (0:42). The Fab’-DNA-Atto647N fluorescence intensity colocalized with the LAT condensate (Fig. 4H), does not increase and decreases in a stepwise fashion as seen in the previous two examples, but changes gradually as Fab’-DNA–TCR complexes dynamically merge and split. Yet, as seen in the previous montages, LAT condensate intensity increases notably after the bound TCR complexes colocalize (Fig. 4I). While not required for T cell activation, these image sequences provide direct observation of long anticipated cooperativity between TCR signaling at the single receptor level.

Consolidation of individual Fab’-DNA–TCR complexes into microclusters is particularly noticeable when T cells first land on SLBs presenting a moderate density (0.1–-0.3 µm^2^) of H57 R97L Fab’-DNA ligand (Fig. 4J) (Movie S8). Single molecules of bound Fab’-DNA are initially well-spaced and there are numerous, small areas of higher LAT density (0:04). Over time, LAT condenses and binding events consolidate until two distinct microclusters of Fab’-DNA–TCR complexes each are colocalized with a large LAT condensate (0:40).

The dynamical nature of the SLB–T cell interface, rapid ligand binding kinetics, and the LAT response significantly complicates systemic quantitative analysis of these described behaviors, and such analysis is beyond the scope of this work. The compilation of observations presented here summarizes notable patterns of cooperativity between binding and LAT condensation that are illuminated by exploring an area of kinetic phase space not available with native ligands.

### High on-rate affects the temporal distribution of cell-wide LAT condensation

We next analyzed cell-wide signaling in response to synthetic Fab’-DNA ligands, focusing on early activation responses (within 5-20 min) to understand how ligand kinetic parameters are initially integrated. As cells landed and spread on SLBs presenting H57 Fab’-DNA and the R97L mutant, these ligands with high on-rates rapidly bound TCR. As a result, many LAT condensates formed concurrently soon after cell landing, and the pace of new LAT condensation subsequently slowed (Fig. 5A and 5B; Movie S9). The number of LAT condensates formed was greater in cells that bind the higher-affinity parental H57 Fab’-DNA compared to the R97L mutant at matched ligand density (0.5 µm^-2^), in line with numerous studies that show higher affinity ligands generate more signaling activity *in vitro* (4–11). As expected, only a few LAT condensates formed in T cells that interacted with bilayers presenting only ICAM. In contrast to the temporal distribution seen for Fab ligands, LAT accumulates linearly in time in cells interacting with pMHC ligands presented at the same density of 0.5 µm^-2^ (Fig. 5C) (13).

**Figure 5.**
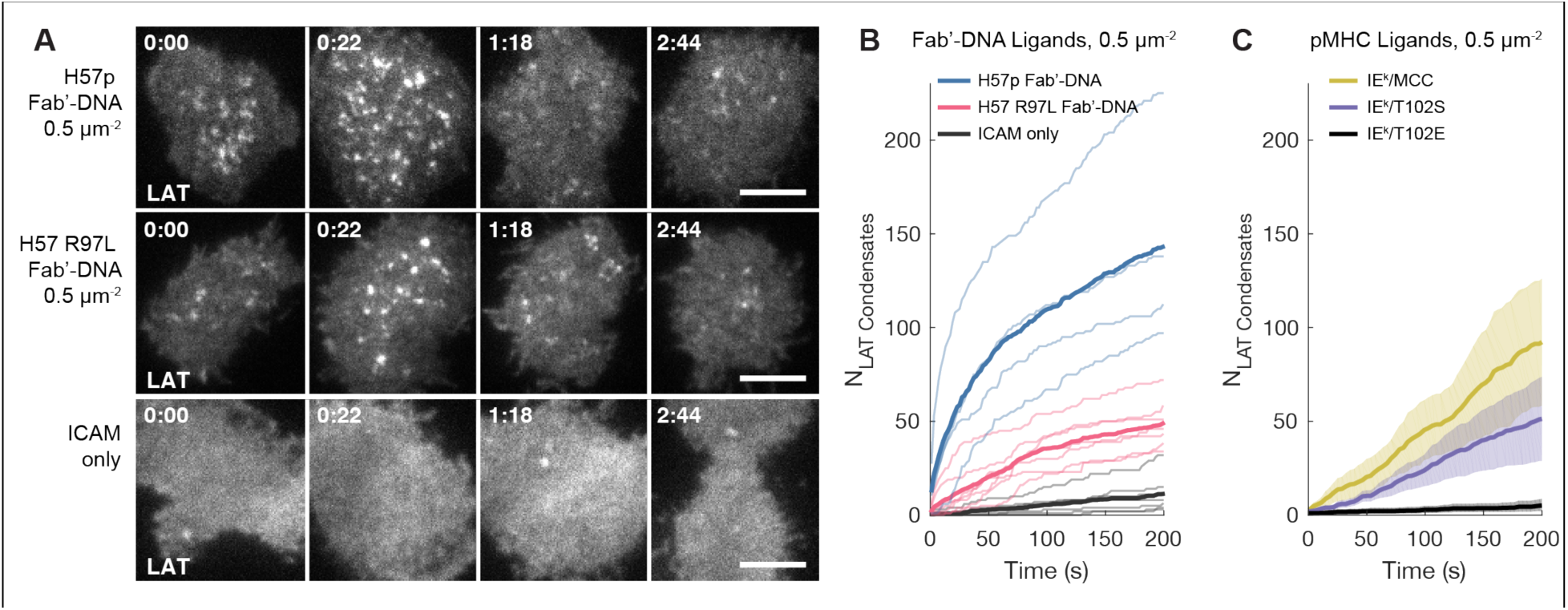
High on-rate affects the temporal distribution of cell-wide LAT condensation. (A) Montages of LAT signal from cells shows a rapid accumulation of LAT condensation soon after landing on Fab’-DNA bilayers, followed by a more moderate rate of LAT condensation accumulation. LAT condensates are sparse in cells on ICAM only bilayers. Scale bar 5 µm. (B) The cumulative number of LAT condensates through time is plotted for cells on bilayers presenting parental, commercial H57 Fab’-DNA (n = 4 cells), H57 R97L Fab’-DNA (n = 7 cells) and ICAM only (n = 7 cells). Individual cell traces are denoted by thin, light lines and the average is denoted by thick, dark lines. (C) The cumulative number of LAT condensates through time is plotted for cells on bilayers presenting pMHC ligands. The mean is plotted as a dark solid line and shaded regions show ± 1 SD. Data are replotted from McAffee et al. 2022.

Other aspects of T cell signaling in response to Fab’-DNA ligand match behaviors typically observed for numerous ligands. H57 R97L Fab’-DNA formed microclusters at the SLB-T cell interface shortly after cell landing when presented at moderate to high densities, and these microclusters co-localized with LAT condensates (Fig. S5A) (55, 56). As cells activate, bound TCR and LAT are shuttled to a kinapse, and central supramolecular activation clusters formed in the immunological synapse in response to high ligand density (Fig S5B) (57). T cell footprints and mobility on bilayers presenting pMHC and Fab ligands are also very similar in the context of ICAM–LFA-1 adhesion (30).

### H57 R97L Fab’-DNA potency falls between MCC and T102S pMHC

Activation of the calcium signaling pathway was determined using a fluorescent NFAT reporter protein. NFAT translocates to the nucleus downstream of calcium flux and is a robust reporter of early T cell activation (36). Unlike calcium flux, which gives an analog response, NFAT translocation can be used to identify cells as either activated or not depending on whether or not fluorescence signal has accumulated in the nucleus. A cell is defined as not activated when the ratio of signal in the nucleus compared to the cytoplasm is less than one (Fig. 6A, left) and active when the ratio is greater than one (Fig. 6A, right). The fraction of activated cells was assessed as a function of bilayer density after cells have interacted with the bilayer for 15 min. The dose-response curve for H57 R97L Fab’-DNA (τ = 4.4 s) falls between that of IE^k^/MCC (τ = 50 s) and IE^k^/T102S (τ = 9 s) (Fig. 6B) (10, 34). H57 R97L Fab’-DNA activates a similar fraction of cells at low antigen densities as the much longer-dwelling IE^k^/MCC, perhaps due to coordinated binding events increasing the size and duration of LAT condensates (as analyzed in Fig. 4) and/or the burst of concurrent LAT condensation seen upon cell landing on bilayers presenting Fab ligands (as shown in Fig. 5). At higher densities, H57 R97L Fab’-DNA more closely resembles the shorter-dwelling IE^k^/T102S. The shallower dose-response curve for H57 R97L Fab’-DNA compared to pMHC ligands could be due to binding kinetics and/or differences in how ligand binding couples to other signaling responses such as co-receptor engagement.

**Figure 6.**
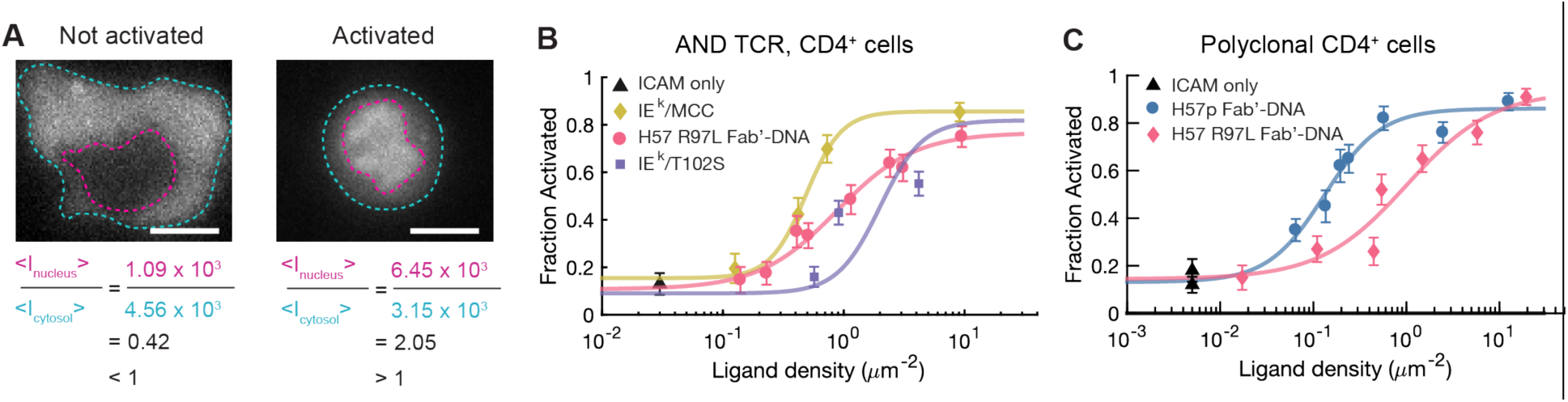
H57 R97L Fab’-DNA activates clonal and polyclonal T cells. (A) Cells are defined as activated if the NFAT-mCherry fluorescent signal is greater in the nucleus compared to the cytosol. Scale bar 5 µm. (B) NFAT dose-response curve for H57 R97L Fab’-DNA in CD4+ T cells expressing the AND TCR falls between IEk/MCC and IEk/T102S. Fits assume that the maximal activation for all ligands is the same. Error denotes SEM. n > 70 cells for each data point. (C) NFAT dose-response curves of polyclonal CD4+ T cells for parental H57 Fab’-DNA and H57 R97L Fab’-DNA show that the R97L mutant is a less potent ligand. Error denotes SEM. n > 100 cells for each data point. Data are representative of three independent experiments.

We assayed IL-2 release after 6 h of exposure to H57 R97L Fab’-DNA bilayers with an enzyme-linked immunosorbent assay (ELISA). Cells exposed to H57 R97L Fab’-DNA produced more IL-2 than cells on bilayers presenting only ICAM, but not nearly as much as cells presented with a high density of IE^k^/MCC or a saturated surface of H57 antibody (Table S1). Multiple iterations of the IL-2 ELISA showed highly variable absolute IL-2 production, and results cannot be interpreted quantitatively. The extent of downstream T cell activation from Fab’-DNA ligands remains an open question worth exploring for the development of universal T cell receptor agonists.

### Polyclonal T cells discriminate Fab’-DNA in a dwell-time-dependent manner

To illustrate the broad applicability of Fab’-DNA constructs, we investigated the activation of effector T cells from polyclonal B6 mice with both parental H57 Fab’-DNA from commercially available antibodies and the lower-affinity expressed H57 R97L Fab’-DNA. The dose-response curve for R97L is shifted significantly to the right compared to parental H57, requiring nearly ten times the amount of ligand present on the bilayer to reach half-maximal activation (1 µm^-2^ compared to 0.15 µm^-2^) (Fig. 6C). The shift to a greater EC50 with decreased ligand dwell time aligns with observations that, *in vitro*, cellular activity correlates well with kinetic off-rate and affinity (4–11). The dose-response curve for both polyclonal (Fig. 6C) and clonal (Fig. 6B) CD4^+^ cells in response to H57 R97L Fab’-DNA are very similar, both with ED50 of about 1 µm^-2^ and exhibiting a shallower slope than seen for pMHC ligands. Polyclonal CD4^+^ cells were isolated from B6 lymph nodes and spleens to best compare to the CD4^+^ AND T cells (Fig. S6A), but experiments conducted with the full population of CD4^+^ and CD8^+^ T cells from B6 mice yielded similar results (Fig. S6B). These results show that, like with pMHC, T cells respond differentially to Fab’-DNA constructs with varied affinity.

## DISCUSSION

### Effector T cells read the duration of single binding events similarly for Fab’-DNA and pMHC ligands

The Fab’-DNA and pMHC ligands differ both in binding epitope and interaction with co-receptor. Thus it was unclear if T cells would read the dwell time of the two classes of ligands in a similar manner. In particular, coreceptor scanning (41), requires the coreceptor docking site on pMHC to efficiently bring active Lck to pMHC–TCR complexes, providing a tunable feature to the TCR ligand discrimination mechanism. In light of this, our observation of essentially identical single TCR antigen discrimination functions for the two ligands may be surprising. However, this is likely a matter of differing timescales and/or cell types. Coreceptor scanning effects are likely more significant near the short dwell time threshold (∼0.5 s dwell times) and for thymocytes compared to effector T cells (40). Even with a mean dwell time of only 4.4 s, the H57 R97L Fab’-DNA ligand is still a moderate agonist and well beyond minimal activation dwell times (40).

### Direct observations of the cooperative interplay between ligand binding and LAT condensation unify models of ligand discrimination

While long, isolated TCR binding events are sufficient to activate T cells (10, 34), cooperativity between TCR has long been speculated in the literature (7, 8, 10, 33, 45, 46). By tracking the interplay between correlated H57 R97L Fab’-DNA–TCR binding events and LAT condensation, we directly visualize several notable cooperative behaviors: (i) Multiple correlated binding events enhance LAT condensation size and duration; (ii) additional Fab’-DNA–TCR binding exhibits a higher on-rate within preexisting LAT condensates; (iii) bound TCRs become more concentrated at condensates, though they can move in and out of the area. These visualizations can help unify interpretation of a wide range of experimental observations as illustrated in the following model of kinetic ligand discrimination.

Fig. 7 depicts a model, informed by distinct behaviors observed with the H57 R97L Fab’-DNA, that emphasizes a role for LAT condensation in coordinating cooperativity among TCRs. As has been extensively characterized, individual binding events trigger TCR through a kinetic proofreading mechanism (58, 59) involving the Src family kinase Lck and Syk family kinase ZAP-70 (12, 60) and exclusion of the phosphatase CD45 (61, 62), which increases the local density of phosphorylated LAT (pLAT) (Fig. 7A). Some of these pools of pLAT may independently condense without input from neighboring TCR, with a probability that depends on ligand dwell time (Fig. 7B)(13). Local pools of pLAT can also serve as a short-term memory of nearby TCR activation and integrate signal across different TCR in the same spatiotemporal neighborhood (Fig. 7C). As with other biomolecular condensates (63), phosphorylation-driven LAT condensation (37, 38) exhibits a switch-like response regulated by controlling the abundance of molecular components relative to the condensation point. Therefore, serial triggering of multiple nearby TCR by a single ligand (43) could enhance the probability of LAT condensation beyond what the dwell times of single TCR binding events predict—primarily by allowing more activated ZAP-70 molecules on several different TCRs to contribute to the same LAT condensation event. The signaling consequences of such serial triggering behavior have been repeatedly observed particularly under experimental conditions with slow ligand mobility (7, 8, 33, 43, 46). With the direct visualization afforded by the unique H57 R97L Fab’-DNA binding kinetics, we identify LAT condensation as the means by which multiple binding events are integrated.

**Figure 7.**
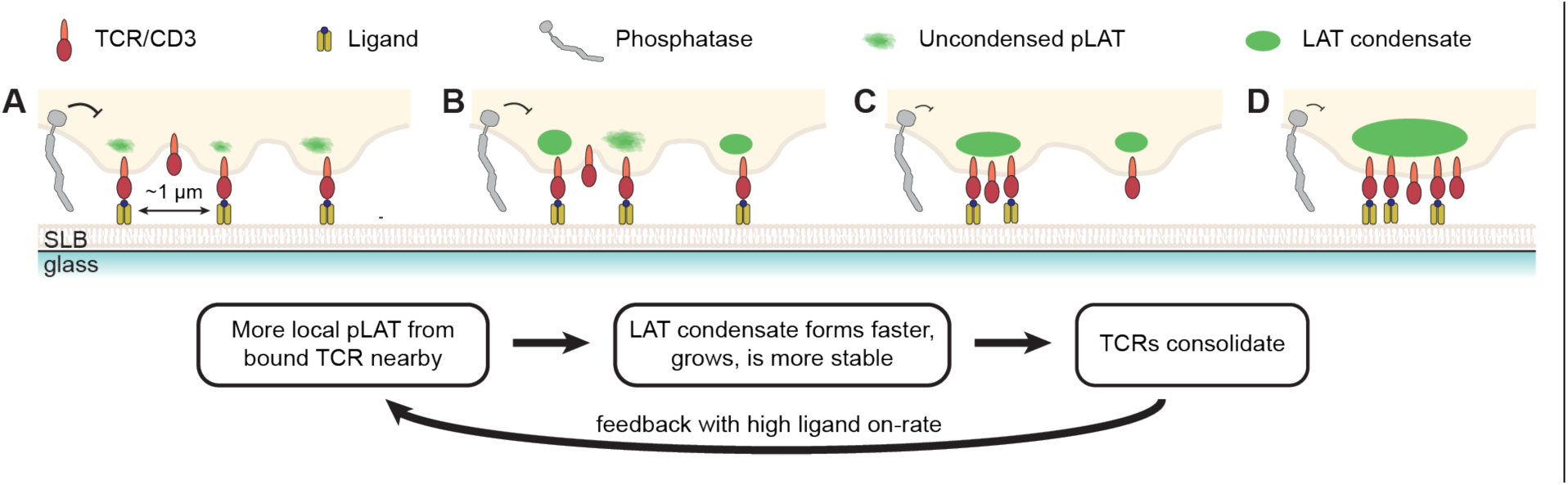
A unified model of kinetic ligand discrimination shows the interplay between ligand binding, LAT condensation, and TCR clustering. (A) Individual ligand–TCR binding events generate signaling activity through ZAP70 recruitment and activation and phosphatase exclusion. Phosphorylated LAT accumulates locally. (B) LAT condenses, either from a single bound TCR or with pLAT from multiple bound TCR. (C) As LAT condenses, TCRs are simultaneously incorporated. The LAT condensate stabilizes with input from multiple bound TCR. (D) The LAT condensate drives a colocalized microcluster of TCR, resulting in a higher local on-rate. More binding events from the higher on-rate sustain the LAT condensate. At later times, negative regulatory processes drive the dissociation of LAT condensates.

Lastly, a positive feedback loop occurs as the condensate enhances the local ligand on-rate and consolidates bound TCRs (Fig. 7D). Higher ligand on-rates have previously been observed in signaling hot spots (64), similar to the higher Fab’-DNA on-rate in LAT condensates that we observe here. The positive feedback model highlights how TCRs and pMHCs can rapidly transition from a purely random distribution (65, 66) to a clustered distribution with high signaling activity (55, 67, 68) when interacting with a high density of high-affinity ligand. In agreement with this experimentally observed behavior, LAT condensates were found to form faster in areas of higher local density of bound TCR in a recent computation model (69). Artificial clustering of ligated TCRs likely bypasses native proofreading mechanisms by stimulating significant local phosphorylation without needing positive feedback from LAT condensation.

### Cell-wide timing of LAT condensation could impact the coordination of cellular activation response

The T cell signaling network involves multiple branches that are coordinated in space and time. Rapid, concurrent LAT condensation, as induced by antibody-based ligands with high on-rates, likely affects the timing and localization of other signaling molecules such as pERK and SHP-1, which could alter cellular fate (2, 70). The temporal change in cell-wide signal accumulation may explain the shallower dose-response curve for Fab’-DNA ligands in comparison to pMHC ligands. At low ligand density, Fab’-DNA ligands provide more opportunities for cooperativity than dwell-time-matched pMHC which may increase the probability of cellular activation. But at higher ligand density, Fab’-DNA appears to lose its advantage. Modulating the timing of cell-wide LAT condensation with high-on-rate ligands could have a significant impact on signaling events further downstream such as cytokine secretion and proliferation.

## MATERIALS AND METHODS

### Cloning the H57 Fab’ into a pCES1 Expression Vector

The pCES1 Fab expression vector (47) was a kind gift from the Charles Craik lab at UCSF. His- and myc-tags are included at the C-terminus of the heavy chain for Fab purification and Western blotting applications. The protein sequence for the H57 Fab heavy chain and light chain were determined from the crystal structure of the H57 Fab bound to TCRβ (49). Light and heavy chain sequences were codon optimized for expression in *E. coli* and ordered from Integrated DNA Techonologies (IDT) (Coralville, IA). Flags of 20-30 amino acids matching the plasmid on either side of the Fab sequence were appended to the H57 heavy chain and light chain sequences for incorporating the gene blocks sequentially with Gibson Assembly (reagents supplied by the QB3 Stanley MacroLab at UC Berkeley). A GGGC linker was appended to the C-terminus of the heavy chain for conjugating the Fab fragment to the fluorescently-labeled DNA oligonucleotide functionalized at the 5’ end with a reactive maleimide group, and a TEV recognition site was later added between the GGGC linker and the His-tag using site-directed mutagenesis (Q5 site-directed mutagenesis kit, New England Biotech, Cat #: E0554S, Ipswich, MA). Cleaving off the His-tag after purification over a Ni^2+^ column prevented multivalent interactions between Fab’-DNA and supported lipid bilayers (SLBs) that are functionalized with the complementary oligonucleotide and Ni^2+^-chelated nitrilotriacetic acid (NTA). Site-directed mutagenesis was also used to introduce point mutations in the complementarity-determining region 3 (CDR3) of the H57 Fab heavy chain in order to lower the Fab binding affinity. Flow cytometry was tested to screen several Fab mutants for decreased binding affinity, but the results were difficult to interpret (see SI Materials and Methods for details). The R97L mutant was chosen for large-scale expression and conjugation to DNA because a similar mutation substantially decreased binding kinetics in the anti-CD3 antibody OKT3 (50). Sequences for gene blocks and primers are shown in SI Appendix Tables S2.

### Fab’ Expression and Purification

Fabs were expressed and purified as described in Kim *et al.* (48). On day 1, BL21-CodonPlus (DE3) competent cells were transformed with the Fab plasmid and plated on Luria broth agar plates containing 100 µg /mL carbenicillin for growth overnight. On day 2, four starter cultures of 50 mL 2xYT, 2% glucose, and 100 µg/mL carbenicillin were inoculated each with one colony. Starter cultures were grown overnight at 30 °C, 200-250 rpm. On day 3, each starter culture was diluted into 1 L of 2xYT, 0.1% glucose, and 100 µg/mL carbenicillin to OD600 of about 0.05. Large cultures were grown at 37 °C at 200-250 rpm to an OD600 of about 0.6. Upon reaching an OD600 of 0.6, large cultures were induced with IPTG to a final concentration of 1 mM and grown overnight strictly at 20 °C and 200 rpm.

Expressed Fab, directed to the oxidizing periplasm by signal sequences on both the heavy and light chains, was isolated from the periplasm on day 4. Bacteria were collected by centrifugation at 5,000 g for 30 min at 4 °C. Each pellet was thoroughly resuspended in 15 mL of ice-cold, freshly prepared TES buffer (0.2 M Tris pH 8, 0.5 mM EDTA, 0.5 M sucrose) and transferred to a 50 mL falcon tube to incubate for at least 1 h on an orbital shaker at 4 °C. Ice cold Milli-Q water was then added, either 15 mL or to an approximate volume of 50 mL total solution, and the tubes were incubated for another 45 min on an orbital shaker at 4 °C. The falcon tubes were then centrifuged at 10,000 g for 30 min at 4 °C and the supernatant was saved as the periplasmic fraction.

Next, Fabs were isolated from the periplasmic fraction using a Ni^2+^ affinity column. To do this, 1mL Ni^2+^ agarose bead (Cat # R90101, ThermoFisher Scientific, Waltham, MA) slurry per liter of cell culture were washed in wash buffer 1 (50 mM Tris pH 8, 250 mM NaCl, filtered) three times, centrifuging at 900 x g for two minutes, removing supernatant, and resuspending. 100 µL of 1 M MgCl_2_ and imidazole to a final concentration of 10 mM were added to each periplasmic fraction, and tubes were gently inverted to mix. Washed Ni^2+^were then added to each fraction and allowed to incubate for 1 h with slow rotation. Beads were then spun down at 2000 x g on a tabletop centrifuge for 10 min at 4 °C. Supernatant was decanted and saved as flowthrough and beads were transferred to a column, using wash buffer 1 to resuspend the beads to aid in the transfer.

In the column, beads were washed with 10-20 column volumes of wash buffer 2 (50 mM Tris pH 8, 500 mM NaCl, 20 mM Imidazole). Elution buffer (50 mM Tris pH 8, 500 mM NaCl, 500 mM Imidazole) was then added gently to the column when the final wash nearly reached the surface of the settled beads, totaling 10 column volumes. The first 500-750 µL of flowthrough was collected as void volume. Two elution fractions of 3 mL each were then collected from the column. Elution fractions were dialyzed overnight in 1 x PBS and 2 mM EDTA using 10 kDa MWCO dialysis cassettes (Cat # 66380, ThermoFisher Scientific, Waltham, MA).

On day 5, the His_6_ tag used for Fab purification was cleaved using TEV protease, obtained from QB3 Stanley Macrolab. First, elution fractions were combined and concentrated to 0.5 – 1 mg/mL using Amicon Ultra-4 30 kDa MWCO centrifugal filter (Cat #: UFC803024, Millipore Sigma, Burlington, MA) TEV protease in 25 mM HEPES buffer, pH 7.5, 400 mM NaCl, 10% glycerol, and 1 mM DTT was added to Fab sample at a 1:22 w/w ratio and incubated at room temperature for 1 h. The reaction was then buffer exchanged over a NAP-5 or NAP-10 desalting column (Cat #: 17085301,17085401, Cytiva, Marlborough, MA) that was equilibrated in wash buffer 1.5 (50 mM Tris pH 8, 250 mM NaCl, 10 mM Imidazole). Ni^2+^beads, about 50 times less than the amount used for binding Fab in the periplasm prep, were also equilibrated in wash buffer 1.5 and then incubated with the buffer exchanged Fab sample. Uncleaved Fab and His-tagged TEV protease were thereby removed from the cleaved Fab, which remained in solution. After incubating the Fab sample with Ni^2+^ beads for 1 h at 4 °C, beads were centrifuged down and the supernatant containing cleaved H57 Fab was saved. Beads were washed twice in wash buffer 2, and supernatants of washes were saved.

Fractions collected along the way during the Fab purification were analyzed by SDS-PAGE. The supernatant from the last Ni^2+^ bead incubation and washes containing cleaved Fab were combined and dialyzed overnight in 1 x PBS and 2 mM EDTA using 10 kDa MWCO dialysis cassettes.

### Fab’-DNA Synthesis

On day 6, the purified Fab was conjugated to the 3’-thiol-fluorophore, 5’-amine-linker DNA oligonucleotide (sequence: 5’-/5AmMC6/GGTGTGATGTATGTGGA/3ThioMC3-D/-3’ ordered from IDT). The synthesis of this functionalized oligonucleotide is described in Lin *et al.* (30). 4-fold excess dye-DNA-linker was added to Fab sample (concentrated to at least 0.4 mg/mL using an Amicon Ultra-4 10 kDa MWCO centrifugal filter) and incubated at room temperature for 3 h, shaking. T remove excess dye-DNA-linker after the reaction, the solution was diluted to up to 4 mL and concentrated through an Amicon Ultra-4 30 kDa MWCO centrifugal filter. Dilution and concentration were repeated, then the solution was moved to a new Amicon Ultra-4 unit and the solution was filtered two more times.

Concentrated Fab’-DNA was then purified over a size exclusion column (Superdex 200 Increase 10/300, Cytiva) using 1 x PBS buffer and an anion exchange column (Mono Q 5/50 GL, Cytiva) using a 20 mM Tris pH 8 with a salt gradient from 150 mM to 1 M NaCl over 40 min and a flow rate of 1 mL/min. Fractions containing purified Fab’-DNA as evaluated by SDS-PAGE were aliquoted and frozen in 10% glycerol.

### Preparing Viral Titer for Transduction

Platinum-Eco (PLAT) viral packaging cells (Cell Biolabs, San Diego, CA) were cultured in PLAT media (Gibco DMEM liquid medium (-sodium bicarbonate, pyruvate, or L-glutamine), heat inactivated FBS, 1 mM Sodium Pyruvate, 100 units/mL Penicilin/Streptomycin, 2 mM L-Glutamine) in a tissue culture-treated T75 flask to reach 50-60% confluency on the day murine organs were harvested (Day 1). 60 µg of linear, polycationic polyethylenimine (Sigma Aldrich) and 15 µg lat-egfp-p2a-nfat-mcherry bicistronic gene in the MCSV backbone (similar to AddGene #91975) were pre-mixed in 1 mL of Opti-MEM media (Life Technologies Cat #:51985-034) and incubated for 20 min. Construction of this plasmid was previously described in ref (13). PLAT cells were transfected by adding this solution to the PLAT media. Media was exchanged to RVC (all components of PLAT media, plus 1 x MEM essential vitamins, 1 x nonessential amino acids, 665 µM L-Arg, 272 µM L-Asn, 14 µM Folic acid, 2 mM L-Glutamine, 50 µM β-Mercaptoethanol) after 4-5 h for collecting viral titer.

### Primary T Cell Harvesting and Culture

All animal work was performed with prior approval by the Lawrence Berkeley National Laboratory Animal Welfare and Research Committee (LBNL’s IACUC), under the approved protocol 177003. T cells were harvested from mice expressing the AND TCR, bred as a cross of B10.Cg-Tg(TcrAND)53Hed/J X B10.BR-H2k2 H2-T18a/SgSnJ from Jackson Laboratory. Polyclonal primary murine T cells were a gift from the Art Weiss lab at UCSF.

Murine spleen and lymph nodes were harvested and processed on Day 1, Organs in ice-cold RVC media were processed through a 70 µm cell strainer, with lymph nodes and spleens treated separately. After centrifuging for 5 min at 500 x g, spleenocytes were resuspended in 5 mL Ack lysis buffer (150 mM NH_4_Cl, 10 mM KHCO_3_, 0.1 mM Na_2_EDTA, pH 7.2–7.4) and pipetted to rupture red blood cells while minimizing bubbles in solution. Spleenocytes were then diluted to 15 mL with RVC, centrifuged, and resuspended in about 4 mL RVC. Lymphocytes and spleenocytes were combined and filtered dropwise through a 40 µm cell strainer, counted, and diluted with RVC to 10-12 million cells/mL. For AND T cells, MCC88-103 peptide (ANERADLIAYLKQATK) was added to the media at a concentration of 2 µM and cells were distributed in a coated 24-well tissue culture plate. Polyclonal cells were plated in 24-well uncoated tissue culture plates that had been treated the night before with 2.5 mg/mL anti-CD3 2C11 and 0.5 mg/mL anti-CD28 antibodies in 1 x PBS, left to incubate overnight, then washed with RVC to introduce a fibronectin coat in the morning.

IL-2 (Sigma Aldrich Cat #: I0523-5X20UG) was added to a final concentration of 1.25 µg/mL on Day 2, within 24 h of organ harvest.

For polyclonal cells, a step was added to isolate CD4^+^ cells in order to more directly compare results from the polyclonal cells to the CD4^+^ AND T cells. On day 3, before transduction with viral titer from the PLAT cells, CD4^+^ cells were isolated using DynabeadsTM UntouchedTM Mouse CD4 Cell Kit (Cat #: 11416D, ThermoFisher Scientific) according to the manufacturer’s instructions. 50-75 million cells were treated for each set of imaging experiments.

T cells used to look at intracellular signaling events were transduced on Day 3 with lat-egfp-p2a-nfat-mcherry packaged in MSCV retrovirus. Cell survival rate on day 3 was usually 60% of the cell count on day 1. Viral titer was removed from PLAT cells and filtered though a 0.22 µm filter to remove cell debris. Cells were centrifuged down at 500 x g for 5 min, exchanged into viral titer at 1.3 million /mL, IL-2 at 1.25 µg/mL, and polybrene at 4 µg/mL (Sigma Aldrich Cat #:TR1003G). They were then distributed to a 24-well plate with 1.5 mL per well, and centrifuged for 1 h at 1330 x g at room temperature. After centrifugation, plates were returned to the incubator. The following day (day 4), cells were exchanged to fresh RVC and IL-2 and recovered at 2.5 – 3.5 million /mL.

Cells were imaged on days 5-8 and were maintained in fresh RVC media containing IL-2. A successful transduction resulted in about 30% of cells expressing the fluorescent reporter proteins.

### Supported Lipid Bilayer Preparation

Low-defect, fluid supported lipid bilayers (SLBs) were prepared as described in (35). Small unilamellar vesicles (SUVs) were first prepared by mixing 95% 1,2-dioleoyl-sn-glycero-3-phosphocholine (DOPC), 3% 1,2-dioleoyl-sn-glycero-3-phos-phoethanolamine-N-[4-(p-maleimidomethyl)cyclohexane-carboxamide] sodium salt (MCC-DOPE), and 2% 1,2-dioleoyl-sn-glycero-3-[(N-(5-amino-1-carboxypentyl)iminodiacetic acid)succinyl] nickel salt (Ni-NTA-DOGS) phospholipids in chloroform in a piranha-etched 25 mL round-bottom flask. Lipids were dried by rotary evaporation and then gently resuspended in Milli-Q water (MilliporeSigma, Billerica, MA) to 0.5 mg/mL total lipid concentration. This solution was sonicated (15 s pulse with 10 s pause for a total of 1 min) with a probe sonicator (Analis Scientific Instruments, Namur, Belgium) in an ice water bath to form SUVs. The SUVs were then centrifuged (21,000 x g, 4 °C, 20 min) to remove titanium particles and lipid aggregates. The supernatant was removed and mixed 1:1 with 1x PBS, resulting in a spreading solution with a final concentration of 0.25 mg/mL total lipid content. SLBs were formed by vesicle fusion of SUVs by adding this spreading solution to Attofluor imaging chambers (Cat #: A7816 by Invitrogen, ThermoFisher Scientific) assembled with freshly piranha-etched glass coverslips (Thomas Scientific, Swedesboro, NJ). Thiol-DNA, complementary to the Fab’-DNA strand (sequence: 5’-/5ThioMC3-D /CCACATACATCACACC/-3’, IDT), was deprotected with 10 mM tris(2-carboxyethyl)phosphine (TCEP) for 90 min to expose free thiol and then incubated on the bilayer at 1 mM in PBS to obtain a density of approximately hundreds of molecules/µm^2^ on the SLB. These bilayers were stable for up to four days at 4 °C. Before experiments, the SLB was charged with 30 mM NiCl_2_ to ensure stable chelation of polyhistidine-tagged ICAM-1 to the NTA-DOGS lipids. Proteins to be coupled to the bilayer were prepared in imaging buffer (20 mM HEPES, 137 mM NaCl, 5 mM KCl, 0.7 mM Na_2_HPO_4_*7H_2_O, 6 mM D-Glucose, 1 mM CaCl_2_*2H_2_O, 2 mM MgCl_2_*6H_2_O, 0.2 mg/mL bovine serum albumin), added to the imaging chamber, and incubated for 35 min. Fab’-DNA incubation concentrations ranged from 50 pM for single-molecule studies to 10 nM for high-density samples. ICAM-1 was incubated at a concentration of 100 nM.

### Microscopy

A Nikon Ti Eclipse motorized inverted microscope (Technical Instruments, Burlingame, CA) was used to collect all images, as previously described in refs (13, 30, 35). Briefly, A laser launch with 488-, 560-, and 640-nm diode lasers (Coherent OBIS, Santa Clara, CA) was aligned into a custom-built fiber launch (Solamere Technology Group Inc., Salt LakeCity, UT). For TIRF imaging, laser illumination was reflected through the appropriate dichroic beam splitter (ZT488/ 647rpc, Z561rdc with ET575LP) to the objective lens [Nikon (1.47, numerical aperture; 100×). RICM and epifluorescent excitation were filtered through a 50/50 beam splitter or band-pass filters (D546/10×, ET470/40×, ET545/30×, and ET620/60×). All emissions were collected through the appropriate emission filters (ET525/50M, ET600/50M, and ET700/75M) and captured on an EM-CCD (iXon 897DU; Andor Inc., SouthWindsor, CT). All filters were from Chroma Technology Corp. (Bellows Falls, VT). All microscope hardware was controlled for image acquisitions using Micromanager software (71).

All live cell imaging was conducted in a heating stage at 37 °C and 5% CO_2_. Imaging chambers equilibrated at 37 °C for 10 min before adding cells. The density of Fab’-DNA-Atto647N on each bilayer was determined prior to adding cells by taking 20 ms exposure time snapshots of at least 15 fields of view at 6 mW laser power at the sample. Laser power was measured at the sample plane with a Newport Power Meter Model 1918-R controller and a 918D-SL-0D2R. Ligand mobility was determined by taking 500 ms exposure time snapshots at a lower laser power of 0.5 mW at the sample. The immobile fraction of ligands was < 0.5% for bilayers used in this study. For live cell imaging, 0.5–2 million T cells were exchanged from RVC to imaging buffer and added dropwise into the open-face chambers.

All Fab’-DNA-Atto647N on the bilayer were resolved using 50 ms exposure time and 6 mW power at the sample. Binding and unbinding to TCR was seen by change in Fab’-DNA mobility in streaming acquisitions. Bound Fab’-DNA was imaged using 500 ms exposure time and 0.5 mW power for data collected in this study unless otherwise noted.

Fraction bound data were collected on cells that had been interacting with a bilayer (0.1 µm^-2^ H57 R97L Fab’-DNA) for 3-7 min. A single set of snapshots was taken per field of view consisting of (1) RICM to visualize cell footprints; (2) 20 ms of 640 nm TIRF with 4 mW at the sample and 1000 gain to visualize all Fab’-DNA; (3) 500 ms of 640 nm TIRF with 4 mW at the sample and 50 gain to visualize bound Fab’-DNA. The short-exposure time snapshot was taken before the long-exposure time snapshot to minimize photobleaching during the acquisition.

Binding events (500 ms, 0.5 mW at sample in 640 TIRF channel) and LAT signal (70-200 ms, 0.2–0.8 mW at sample depending on LAT-eGFP expression level, in 488 TIRF channel) were sequentially imaged using a multidimensional acquisition with a 2 s time lapse on cells upon initial landing on the SLB until about 5 min after landing. For tracking single binding events and proximal LAT condensation, Fab’-DNA was presented on bilayers at 0.05–0.15 µm^-2^. In addition to monitoring the LAT condensation response, dwell time and on-rate were analyzed using these acquisitions. For imaging global cellular LAT response, Fab’-DNA was presented at 0.5 µm^-2^. Photobleaching for the dwell time distribution bleach correction was assessed using the same imaging parameters as for dwell time data on immobile (DPPC replacing DOPC lipids) SLBs presenting about 0.2 µm^-2^ H57 R97L Fab’-DNA. The immobile bilayers allow single Fab’-DNA molecules to be tracked with high fidelity through time for assessing the photobleaching rate.

Snapshots for NFAT dose-response curves were taken of live cells 15–30 min after initially adding cells to the bilayer. Within this range of time, T cells have had ample time to land on the bilayer and respond to the presented ligand (10), but have not significantly dissociated from the bilayer after activation. Transduced cells were identified by their LAT signal in the 488 TIRF channel and were then imaged with a set of snapshots of RICM, 640 TIRF (bound Fab-DNA-Atto647N), 488 TIRF (LAT-eGFP), and 561 epifluorescence (NFAT-mCherry). The RICM channel was used to identify cells that were adhered to the SLB to ensure that only cells that interacted with the SLB and presented Fab’-DNA ligands were analyzed. NFAT-mCherry was imaged using 561 epifluorescence at three z locations, 0 µm, 3 µm, and 6 µm above the TIRF plane in order to resolve a clear image of the nucleus.

### IL-2 ELISA

30 min prior to seeding cells on bilayers, cells were exchanged to IL-2-free media. 2 million cells exchanged to imaging buffer immediately prior to seeding were confronted with SLBs presenting ICAM and additional Fab’-DNA or pMHC ligands or α-CD3-coated coverglass for 6 h. Ligand density was determined by TIRF prior to adding cells, and cell landing was imaged using RICM to confirm interaction with the substrate. After 6 h, 350 µL of the 1 mL total volume in the imaging chamber was gently siphoned and centrifuged down (5 min at 2,000 x g) to remove cell debris. The supernatant was used for triplicate samples in the ELISA (Biolegend, San Diego CA), carried out according to manufacturers instructions.

### Tracking pMHC and LAT

Fab’-DNA:TCR complexes and LAT condensates were identified and tracked using TrackMate (72). Bound Fab’-DNA identified using the Difference of Gaussian detector, a 0.4 um diameter, 800 threshold, and no median filter. Binding events were tracked with a maximum linking distance of 0.7 µm and skipped frames were not allowed. Each binding event required manual inspection to eliminate occasional tracking and/or detection artifacts. In particular, TrackMate does not take particle intensity into account, so additional localizations needed to be added if multiple bound Fab’-DNA merged to the same diffraction-limited spot. For analyzing single, isolated binding events, such as for Figs. 2 and 3, binding events were considered valid if they did not come within 300 nm of another bound Fab’-DNA.

LAT condensates were highlighted by subtracting a copy of the image that was Gaussian blurred with a radius of 4. Condensates were then identified in TrackMate using the Laurentian of Gaussian detector with a radius of 0.7 um and threshold of 1000, with a median filter. Condensate tracks were linked using a maximum distance of 0.7 µm without allowing skipped frames. All LAT condensate tracks were manually inspected to eliminate occasional detection and/or tracking artifacts. Tracks of at least 4 consecutive frames were considered to be condensates. A Fab’-DNA binding event was considered productive if a LAT condensate trajectory began within 300 nm of the binding event and unproductive if this did not occur.

### Estimated Fab’-DNA On-Rate

The cellular on-rate of Fab’-DNA with the presence of photobleaching was estimated using the same method as in ref (13). Briefly, ligand binding is modeled with the simple biomolecular association reaction:

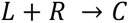

with Fab’-DNA ligand, L, T-cell receptor, R, and complex, C. With receptor density, ᓂ_R_(*t*) and density of *visible,* unbound ligand 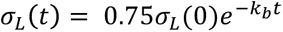, the time-dependent change in density of visible Fab’-DNA:TCR complexes, 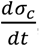, can be estimated as follows:

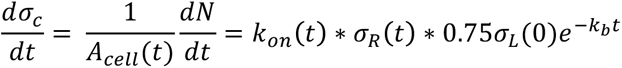

Here, A_cell_ is the area of the cell footprint, N is the total number of visible binding events observed over time t, and k_b_ is the ligand photobleaching rate. The factor of 0.75 is measured empirically as the fraction of unbound ligand across the bilayer. The reaction can be assumed to be at a steady state due to the fast kinetics of H57 R97L Fab’-DNA relative to cellular activation. Absorbing the mean cell area and TCR density into a pseudo kinetic on-rate 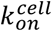 and integrating over time gives:

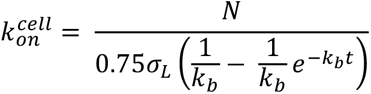

For a collection of 12 T cells across 2 mice, the initial ligand density (σ_L_(0)), total number of binding events (N), and bleaching rate (k_b_), were all determined. The resulting cellular pseudo kinetic on-rate on of H57 R97L Fab’-DNA was found to be 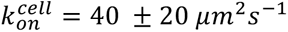.

With this on-rate, we can calculate the expected number of new (visible and bleached) binding events in a spatiotemporal window as:

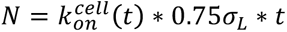

For a ligand density of 0.1 µm^−2^ and an on-rate of 40±20 µm^2^s^-1^, one can expect 180 ± 60 binding events from Fab’-DNA in one cell over one minute. Given a cell footprint of 100 ± 20 µm^2^ from the 12 cells used to calculate the on-rate, this yields 2 ± 1 binding events per µm^2^ contact area per minute. Note that this rate constant is calculated under a low ligand density of 0.1 µm^−2^ and will lose applicability under saturating ligand densities.

### Fraction Bound

The RICM channel was used to crop images to single cells and mask images of all and bound Fab’-DNA molecules. Molecules were localized using TrackMate, and the number of bound Fab’-DNA was divided by the total number of Fab’-DNA for each cell to measure the fraction bound. Cells with fewer than 10 Fab’-DNA under their footprint or for which individual Fab’-DNA molecules were not clearly resolved were excluded from analysis.

### Dwell Time Distribution

Dwell time distributions were compiled from tracks of Fab’-DNA:TCR complexes that remained well-isolated for their entire lifetime. Fluorescent spots resolved for only one frame were discarded from analysis due to occasional spurious localization errors. The observed dwell time distribution, f_obs_(t), was built and fit to a single exponential decay using a custom Matlab script. Reported dwell times were corrected for photobleaching using the following equation:

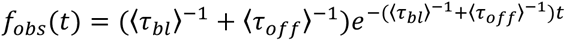

where 〈τ*_bl_*〉 ^-1^ is the fluorophore bleaching rate and 〈τ*_off_*〉 ^-1^ is the ligand off-rate. The bleaching rate was determined by tracking single molecules of Fab’-DNA-Atto647N immobilized in gel-phase bilayers in TrackMate using the same parameters as used for binding events and compiling the survival distribution of these tracks. The rate of photobleaching as a function of frame number was calibrated to time by assuming the same time interval between frames as used for the live cell acquisitions.

### Estimated the Number of Molecules in a LAT condensate

The number of LAT-eGFP molecules within a LAT condensate was calculated as previously described in McAffee *et al*. (13). Briefly, the mean integrated intensity of a single molecules of LAT-eGFP in the plasma membrane was measured from a LAT-expressing cell that was photobleached down to single-molecule densities. The total LAT-eGFP condensate fluorescence intensity was then divided by the intensity of a single molecule, adjusted for differing image acquisition parameters, to calculate the number of fluorescent LAT molecules.

### LAT Success Probability Function and Propensity Function

LAT success probability was calculated by dividing the number of binding events that successfully produced a LAT condensate by the total number of binding events. Progressively larger bins were chosen to minimize noise as the number of events per time window decreased. Data points were fit by a regularized gamma function with a maximum amplitude parameter, c.

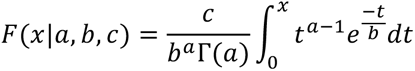

The propensity function (k_c_(t)) is the momentary probability per unit time of a LAT condensate stochastically forming at a specified delay after initial ligand binding. Also known as the hazard rate, it gives the probability of an event occurring given that it has not already occurred. This function is related to the experimentally measured success probability of LAT condensate formation, P_LAT_(t) by the following:

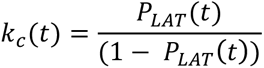

### LAT Delay Time Distribution

The LAT delay time distribution is a direct experimental measure of the propensity function, but it is also convoluted with ligand photobleaching. To build this distribution, the time between the start of a binding event and the start of the associated LAT condensate was histogrammed for all productive, single binding events. For more information, see ref (13).

### NFAT Translocation

A cell was determined to be activated if it contained a greater mean fluorescence intensity in the nucleus compared to the cytosol. Epifluorescent images at 3 µm above the SLB surface were segmented into cytosol and nucleus and the mean intensity ratio of the segments was determined. All images were manually inspected. If the boundary of the nucleus was difficult to determine, the cell was excluded from analysis. All experiments were performed with three biological replicates, and > 70 cells were analyzed per condition. Data were fit by a Hill function with an invariant maximal activation.

### Quantification and Statistical Analysis

Data expressed as x ± y (SD) represents a mean of x and standard deviation of y. SEM refers to y as the standard error of the mean, and (CI) refers to y as the 95% confidence interval. The meaning of the sample size, n, for each data set is specified in relevant figure legends. Quantification and statistical analysis were done using Matlab.

### Data and Code Availability

All raw data and custom analysis scripts and functions (written in Matlab) are available upon request.

## ACKNOWLEDGEMENTS

A community of researchers made this work possible. Many thanks to Kristin Wucherer (Scribe Therapeutics), Koli Basu (Frontier Medicines), and Charles Craik (UCSF) for gifting the Fab expression vector and helping implement Fab expression in the Groves lab; to Serena Meratcioğlu (Vanderbilt University) for aiding in screening Fab mutants; to Lin Shen (University of Chicago) and Arthur Weiss (UCSF) for gifting polyclonal cells from wild type B6 mice; to Scott Hansen (University of Oregon) for synthesizing the LAT-P2A-NFAT plasmid; to Nicole C. Fey (Nutcracker Therapeutics) and Katherine N. Alfieri (UCSF) for expression and purification of pMHCII and ICAM-1, and to members of the Groves lab for many valuable discussions that guided data collection and analysis. The financial support for this work was provided by National Institutes of Health grant P01 AI091580 and by the Novo Nordisk Foundation Challenge Programme as part of the Center for Geometrically Engineered Cellular Systems (NNF17OC0028176).

## AUTHOR CONTRIBUTIONS

KBW designed the research, performed the research, contributed new reagents and analytical tools, analyzed all data, and wrote the paper. AV performed research. JTG supervised research design and data analysis and edited the paper.

