## Supplemental Information for "Differential Roles of Kinetic On- and Off-Rates in T-Cell Receptor Signal Integration Revealed with a Modified Fab’-DNA Ligand"

### Contents

Supplementary Figures 1-6
Captions for Supplementary Movies 1-9
Supplementary Materials and Methods
Supplementary Tables 1 and 2

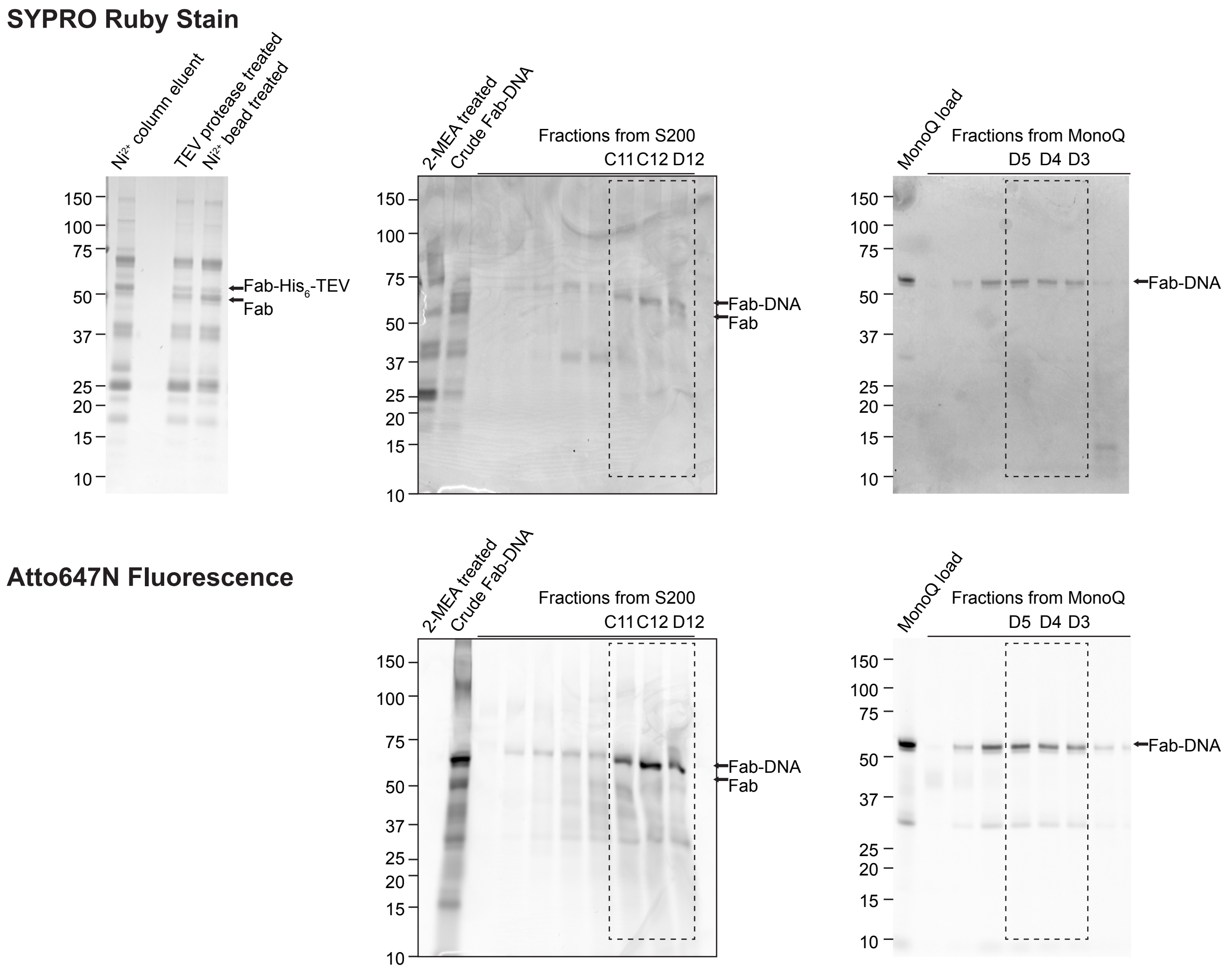


***Figure S1. Synthesis and purification of H57 R97L Fab'-DNA was monitored using SDS-PAGE.*** *(Left) Fab extracted from the periplasm over a nickel column (55 kDa) was treated with TEV protease and uncleaved Fab was removed with additional nickel beads. (Middle) Crude Fab'-DNA was synthesized by conjugating maleimide-functionalized DNA to reduced Fab'. The product was purified by size exclusion and (Right) ion exchange chromotography. Purity was assessed using both SYPRO Ruby staining to visualize all protein and Atto647N fluorescence to visualize the dye-labeled DNA.*

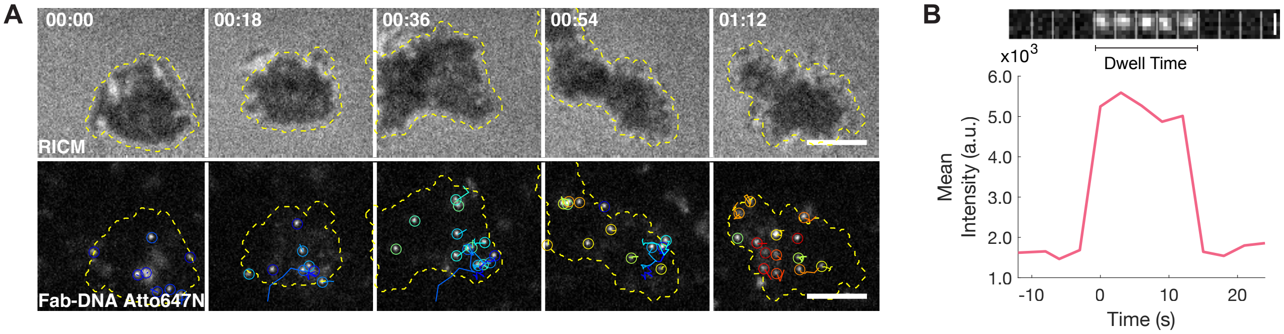


***Figure S2. Measuring in situ ligand dwell times.*** *(A) The cell footprint at the T cell - supported lipid bilayer interface is imaged using RICM, and H57 R97L Fab'-DNA bound to TCR are imaged using TIRF micrsocopy with a long, 500 ms exposure time. Single binding events are tracked through time using TrackMate. Scale bar 5 µm. (E) A single Fab'-DNA molecule binds TCR at time 0 and unbinds or photobleaches 12 s later. The intensity trace of the particle increases and decreases each in a single step, confirming the presence of a single fluorophore. Tracks of single binding events are used to build a dwell size distribution. Scale bar 0.5 µm.*

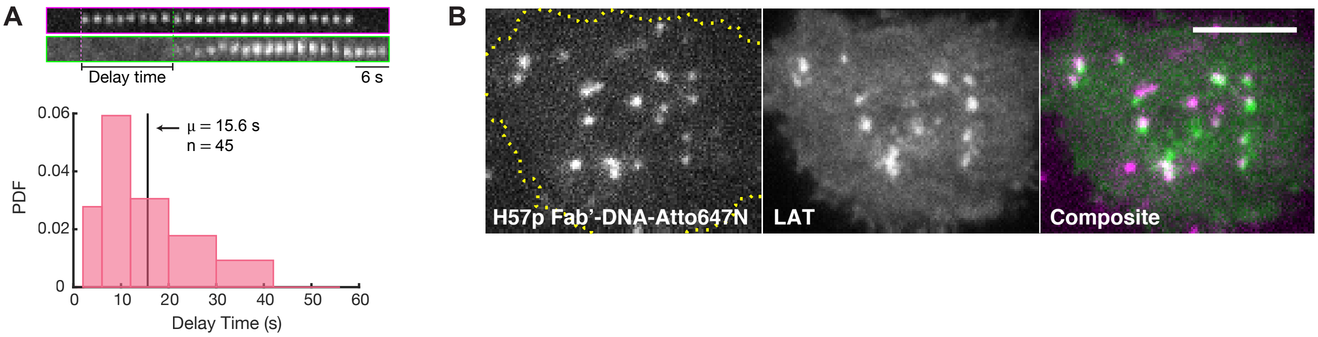


***Figure S3. Activity of isolated Fab'-DNA binding events.*** *(A) The mean delay time between initial H57 R97L Fab'-DNA binding and LAT condensation is 15.6 s. This is significantly longer than the mean ligand dwell time of 4.4 s and illustrates that only rare, long-dwelling Fab'-DNA binding events are productive in isolation. Only binding events that remained isolated for their duration were included in analysis. (B) Single molecules of expressed, parental H57 Fab'-DNA trigger local LAT condensation. Scale bar 5 µm.*

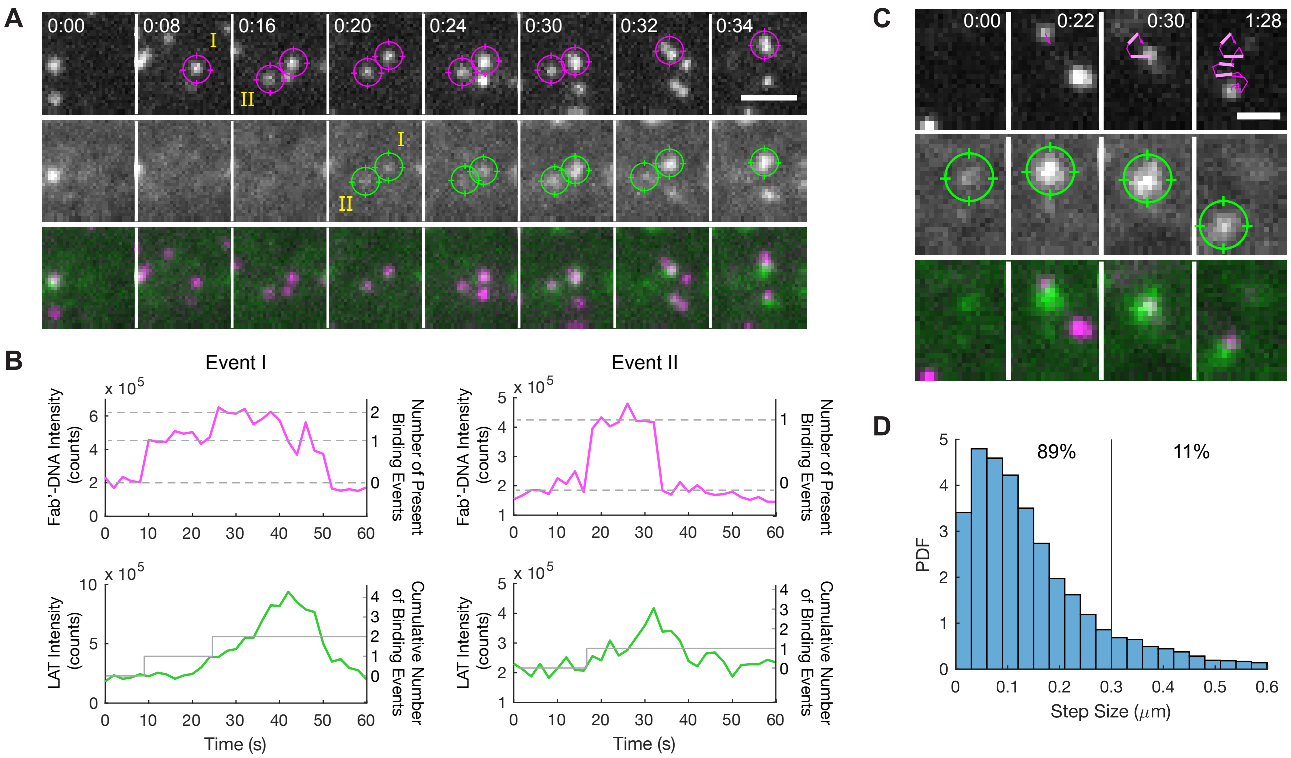


***Figure S4. Additional examples of the interplay between binding events and LAT condensation.*** *(A) Two neighboring binding events both initiate LAT condensation (0:20). The top Fab'-DNA is joined by a second (0:24) and the local LAT condensate grows (0:34), whereas the bottom Fab'-DNA remains isolated from other binding events and the local LAT condensate remains small and short-lived. (B) Fluoresecence intensity traces for binding events (top) and LAT condensates (bottom) for events I (left) and II (right). (C) A molecule of Fab'-DNA binds into an existing LAT condensate. The LAT condensate grows. The large steps (> 0.3 μm; bold light pink) that the binding event takes between some adjacent frames may be due to ligand unbinding and rebinding a different TCR in the area of the LAT condensate. (D) These steps are at the tail end of the step size distribution taken from all binding events tracked with this imaging condition.*

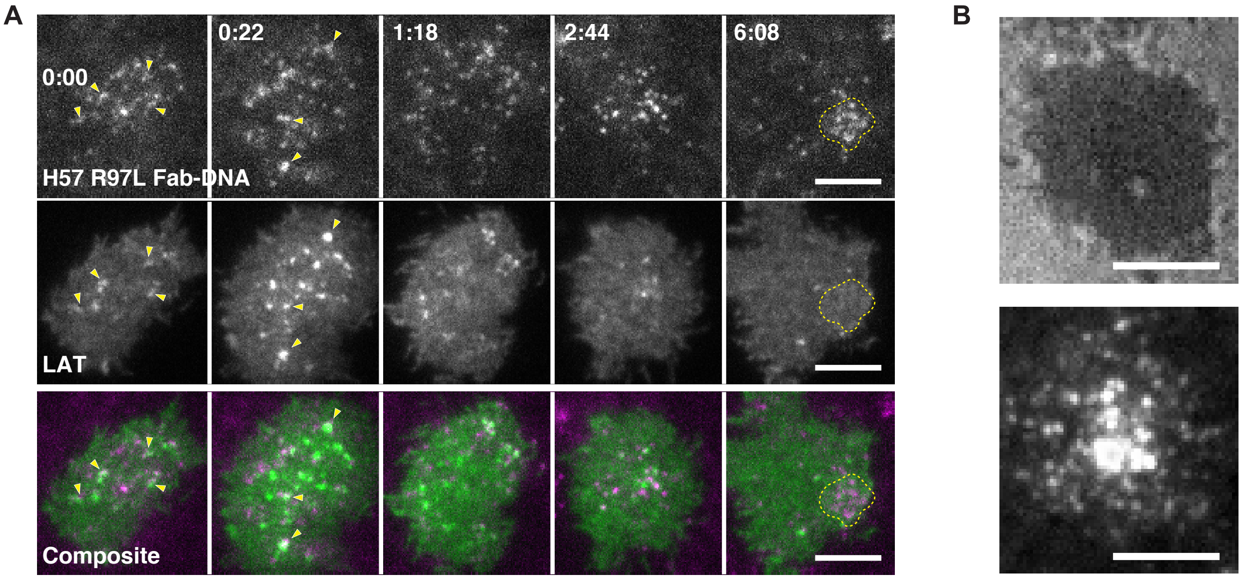


*F**igure S5.** (A) Binding events (top), LAT condensation (middle), and composite (bottom) montages of a cell landing and spreading on a bilayer presenting 0.5 μm$^{-2}$ H57 R97L Fab'-DNA. Yellow arrows indicate examples of colocalization between binding events and LAT condensates. Binding events are directed to a central region of the immmunological synapse (dotted yellow line), as expected, over the course of minutes. LAT frames are duplicated from Fig. 5. Scale bar 5 μm. (B) cSMAC formation in a cell 10 min after landing on a bilayer presenting 6 μm$^{-2}$ H57 R97L Fab'-DNA. Cell footprint (top) and binding events (bottom). Scale bars 5 μm.*

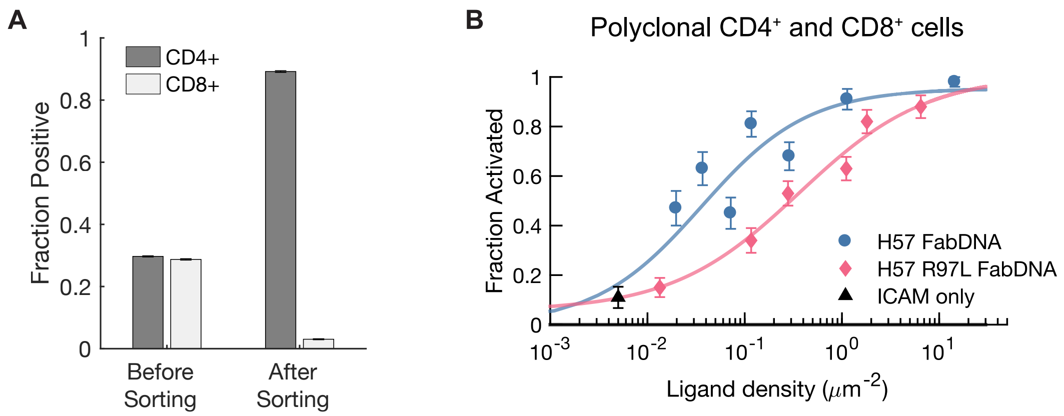


***Figure S6. Sorting polyclonal cells for co-receptor expression does not affect the dose-response curve.*** *(A) CD4 positive cells were sorted from polyclonal, wild type B6 T cells on Day 3 since harvesting. The fraction of cells expressing CD4 and CD8 coreceptor before and after sorting were assessed using fluorescent antibodies. (B) The dose-response curve of unsorted polyclonal cells shows parental H57 Fab'-DNA to be a more potent ligand than H57 R97L Fab'-DNA.*

**CAPTIONS FOR SUPPLEMENTARY MOVIES**

**Movie S1: Single molecules of H57 R97L Fab'-DNA can rapidly rebind to TCRs.** All Fab'-DNA molecules in the bilayer are imaged continuously with 50 ms exposure time to track both bound and unbound ligand. The ligand in box 1 unbinds one TCR at 1.35 s and rebinds a new TCR at 1.45 s. The ligand in box 2 initially binds at 0.55 s, unbinds at 1.70 s, and rebinds at 1.85 s. A montage of box 2 is presented in Fig. 2B. The rapid rebinding of Fabs to new TCRs is indicative of its high on-rate. The outline of the cell footprint is indicated by the dotted yellow line. Scale bar is 5 μm.

**Movie S2: LAT condenses in response to long, isolated H57 R97L Fab'-DNA–TCR binding events.** Bound H57 R97L Fab'-DNA-Atto647N (left, magenta) and LAT-eGFP (middle, green) are imaged in TIRF with a 2 s time interval between frames. Single molecules of Fab'-DNA bind and unbind TCRs at the T cell–SLB interface. LAT condenses in response to single binding events. The composite of both channels is shown on the right. H57 R97L Fab'-DNA is presented at 0.05 $\mu m^{-2}$ on this bilayer. Scale bar 5 μm.

**Movie S3: LAT condensate grows in response to local Fab'-DNA binding.** A single Fab'-DNA ligand (magenta) binds TCR at the SLB–T cell interface and nucleates a LAT condensate (green). Additional Fab'-DNA ligands bind into the area of the LAT condensate. The LAT condensate subsequently grows after new binding events. Scale bar 2 μm.

**Movie S4: A LAT condensate with multiple bound TCR persists while a condensate from a single binding event quickly fades.** Two distinct binding events (magenta) present at the 0:20 time point each nucleate a LAT condensate (green). A new Fab'-DNA ligand binds into the upper LAT condensate at 0:24 and the condensate grows. The lower LAT condensate does not experience additional ligand binding and quickly fades. Other LAT condensates and colocalized bound TCR are present in the field of view as well. Scale bar 2 μm.

**Movie S5: LAT condensation promotes local binding.** Numerous independent Fab'-DNA ligands (magenta) bind to and unbind from TCR in a LAT condensate (green). The LAT condensate grows in response to additional binding events. Scale bar 2 μm.

**Movie S6: A single Fab'-DNA ligand hops within a LAT condensate.** A molecule of Fab'-DNA (magenta) binds into an existing LAT condensate (green) and the LAT condensate subsequently grows. That molecule frequently takes large steps between frames imaged at a 2 s time interval, though it stays within the area of the condensate. At least some of these large steps are attributed to unbinding and rebinding to a new TCR. Scale bar 2 μm.

**Movie S7: Bound TCRs consolidate within LAT condensates but still dynamically exchange.** Two separated Fab'-DNA–TCR complexes (top-right at 0:18 time point; magenta) merge into the same diffraction-limited area (0:24) and stay colocalized for several frames. A local LAT condensate forms, fluctuates for a few frames, then grows with time (green). One of the bound TCRs moves away from the area of the condensate while the other tracks with the condensate. The dynamic exchange of bound Fab'-DNA in and out of the areas of LAT condensates is seen in other areas of the field of view as well. Scale bar 2 μm.

**Movie S8: Interplay between ligand binding, LAT condensation, and TCR clustering.** Soon after a cell lands on a supported bilayer presenting 0.1 $\mu m^{-2}$ H57 R97L Fab'-DNA, LAT condensates are already present. In a central region, outlined in yellow, the transition from single binding events with a local enhanced LAT density to colocalized LAT condensates and

TCR microclusters is particularly obvious. Other examples of the types of the interplay discussed in the text occur across the SLB–T cell interface. LAT condensates typically form in the periphery of the interface and disassemble as they are tracked to the center of the interface, as is seen with pMHC ligands. The region outlined in yellow is presented in Fig. 4J. Scale bar 2 µm.

**Movie S9: LAT condenses quickly and concurrently in T cells confronted with high ligand density (0.5 µm$^{-2}$).** As the cell is landing and spreading, it rapidly forms LAT condensates that transiently correlate with bound TCR. LAT condensates dissociate as they are tracked toward the center of the SLB–T cell interface, and the rate of new LAT condensate nucleation notably slows after about 1 min. Bound Fab'-DNA are tracked to the kinapse over the course of minutes as expected, though they rapidly exchange due to their fast binding kinetics. Scale bar 5 µm.

### SUPPLEMENTARY MATERIALS AND METHODS

#### *Flow Cytometry Screening of Fab Fragments*

A flow cytometry screen of crude periplasm extract from three Fabs with point mutations in CDR3 (R97L, R97A, H100A) showed that all mutants tested had decreased binding affinity to primary murine T cells compared to the parental Fab, but none had a signal that rose significantly above the negative background controls.

Crude periplasm extract was used to test expressed Fab binding affinity to primary murine T cells. H57 Fabs with R97L, R97A, and H100A were tested. One million T cells were incubated in 100 µL crude periplasm extract from 5 mL cultures for 20 min on ice. After washing with imaging buffer, cells were incubated in 100 µL of 5 µg/mL anti-c-myc-AlexaFluor488 antibody, clone 9E10 (Cat #: MCA2200A488, BioRad, Hercules, CA) for 15 min on ice. Cells were washed 3 times, resuspended in 500 µL imaging buffer, and transferred to a flow cytometry tube. The parental H57 Fab and anti-CD4-AlexaFluor488 antibodies (Cat #: 100423, Biolegend, San Diego, CA) were used as positive controls and an irrelevant Fab and no Fab (only buffer) were used as negative controls. Immediately after washing the anti-myc antibody, labeling efficiency was assessed using flow cytometry. The two-step protocol of incubating cells first in the Fab and second in the fluorescent label led to overall weak but detectable signal, comparing the expressed parental H57 Fab to the directly labeled anti-CD4 antibody. The results were difficult to interpret but suggested all of these mutants decreased Fab affinity. Future work should append a fluorescent protein directly to the Fab heavy chain C-terminus in order to minimize the time and number of wash steps required between Fab incubation with T cells and the flow cytometry screen.

**Table S1:** IL-2 ELISA from 1.5 million AND T cells exposed to substrates for 6 h. Error denotes standard deviation from three technical replicates.

| Ligand | IL-2 (pg/mL) | Density ($\mu m^{-2}$) | Range, ≥ 3 experiments* |
|---|---|---|---|
| ICAM | 0.7 ± 0.2 | -- | 0.5 – 2.2 |
| ICAM, H57 R97L Fab'-DNA | 1.74 ± 0.06 | 8.5 | -- |
| ICAM, H57 Fab'-DNA | 0.9 ± 0.1 | 7.0 | 0.9 – 48 |
| ICAM, MCC | 268 ± 5 | 200 | 4.2 – 350 |
| Antibody on glass | 1190 ± 40 | Saturated | 700 – 5000 |

* Data reproduced from Lin JJ *et al.* (2020) *Biophys. J.* 118:1–15.

**Table S2:** Sequences for gene blocks and primers

| | | |
|---|---|---|
| **H57 Fab Light Chain** | GTTCCTTTCTATTCTCACAGTGCACTTTATGAGTTGATTCAGCCAAGTTCCGCC<br>TCAGTAACGGTCGGTGAAACCGTAAAGATCACATGCTCGGGGGACCAATTAC<br>CCAAGAACTTCGCCTATTGGTTTCAGCAGAAGTCGGACAAAAACATTTTATTAT<br>TGATCTACATGGACAACAAGCGCCCAAGCGGCATCCCGGAACGCTTCTCTGG<br>GTCTACGAGCGGTACTACGGCGACATTAACGATCTCAGGAGCTCAACCAGAA<br>GATGAGGCGGCGTACTATTGTCTTAGTTCATACGGTGACAACAATGACTTAGT<br>CTTTGGTAGCGGCACGCAACTTACAGTGCTTCGCGGTCCTAAGTCAAGCCCC<br>AAAGTCACAGTGTTCCCGCCATCCCCGGAAGAGTTGCGCACTAATAAAGCGA<br>CATTAGTATGCTTAGTCAACGACTTTTATCCAGGCAGCGCAACAGTCACCTGG<br>AAGGCAAATGGCGCGACTATTAACGATGGCGTTAAGACCACGAAACCGTCGA<br>AACAGGGGCAAAACTACATGACTTCCTCCTACCTTTCTCTTACAGCGGACCAA<br>TGGAAGAGTCATAACCGCGTGAGTTGCCAGGTCACACATGAGGGCGAAACTG<br>TCGAAAAATCCCTGTCGCCAGCCGAGTGCTTATAATAAGGCGCGCCAATTCTA<br>TTTCAAGGAG | |
| **H57 Fab Heavy Chain** | CAGCCGGCCATGGCCCAGGTGCAGCTGCAGGAAGTGTATTTGGTGGAGTCT<br>GGGGGCGACCTGGTACAACCTGGGAGTAGTTTAAAAGTATCCTGCGCGGCTA<br>GTGGTTTTACTTTCAGCGATTTCTGGATGTATTGGGTTCGCCAGGCACCCGGA<br>AAGGGTCTGGAATGGGTCGGCCGCATTAAAAACATCCCCAATAATTACGCGA<br>CTGAATATGCGGACTCGGTGCGTGGTCGCTTCACGATTAGCCGTGACGATTC<br>ACGTAATAGTATCTACTTGCAAATGAATCGTTTACGTGTTGACGACACCGCAAT<br>TTACTATTGCACCCGTGCGGGCCGCTTTGACCACTTTGACTACTGGGGTCAAG<br>GAACGATGGTCACTGTCTCTTCGGCAACCACAACGGCACCTTCCGTCTATCCC<br>TTAGCACCTGCCTGCGACAGTACAACTTCGACGACCGACACTGTTACATTGGG<br>CTGTTTAGTCAAAGGATACTTCCCAGAACCCGTTACAGTAAGTTGGAACAGTG<br>GCGCATTGACGTCTGGAGTCCATACGTTTCCATCGGTGCTTCACTCTGGGCTT<br>TATAGCCTGTCCTCTAGTGTAACTGTCCCATCCTCAACATGGCCTAAGCAGCC<br>GATTACATGTAATGTGGCTCATCCGGCGTCGTCTACAAAGGTGGATAAGAAAA<br>TTGAACCGCGTGGGGGTGGTTGTGCGGCCGCACATCATCATCACCATCACGG<br>GGCCGCA | |
| | Forward | Reverse |
| **Add TEV** | cttccagtcaCATCATCATCACCATCACGG<br>GGCC | tacaggttttcTGCGGCCGCACAACCACC |
| **R97L** | CGTGCGGGCCtCTTTGACCAC | GGTGCAATAGTAAATTGCGGTG |
| **R97A** | CCGTGCGGGCgcCTTTGACCAC | GTGCAATAGTAAATTGCGG |
| **H100A** | CCGCTTTGACgcCTTTGACTACTGGG<br>GTCAAG | CCCGCACGGGTGCAATAG |
